# The telomere regulatory gene POT1 responds to stress and predicts performance in nature: implications for telomeres and life history evolution

**DOI:** 10.1101/2021.01.06.425609

**Authors:** Sarah E. Wolf, Tiana L. Sanders, Sol E. Beltran, Kimberly A. Rosvall

## Abstract

Long telomeres have become nearly synonymous with a variety of fitness-related traits and may be mediators of ecologically relevant variation in life history strategies. Growing evidence suggests that telomere dynamics are more predictive of performance than length itself, but very little work considers how telomere regulatory mechanisms respond to environmental challenges or influence performance in nature. Here, we combine observational and experimental datasets from free-living tree swallows (*Tachycineta bicolor*) to assess how performance is predicted by the telomere regulatory gene POT1, which encodes a shelterin protein that sterically blocks telomerase from repairing the telomere. First, we show that lower POT1 gene expression was associated with higher female quality, i.e. earlier breeding, and heavier body mass. We next challenged mothers with an immune stressor (lipopolysaccharide injection) that led to ‘sickness’ in mothers and 24h of food restriction in their offspring. While POT1 did not respond to maternal injection, females with lower constitutive gene expression were better able to maintain feeding rates following treatment. Maternal injection also generated a one-day stressor for chicks, who responded with decreased POT1 gene expression and elongated telomeres. Other putatively stress-responsive mechanisms (i.e. glucocorticoids, antioxidants) were not significantly different between control and stress-exposed chicks. Model comparisons indicated that POT1 mRNA abundance was a largely better predictor of performance than telomere dynamics, indicating that telomere regulators may be powerful modulators of variation in life history strategies.

## 1. INTRODUCTION

Ecology and evolutionary biology seek to understand causes and consequences of variation in life history strategies. In the last 30 years, biomedical research has revealed telomeres as strong predictors of health and longevity, and integration of these perspectives into eco-evolutionary biology points to telomeres as important mediators of life history trade-offs. Telomeres are ribonucleoprotein structures that buffer chromosomes from erosion during cellular replication (Zakian, 2012) but consequently shorten over time (Allsopp et al., 1995; De Lange, 2009), especially during periods of rapid growth (Geiger et al., 2012; Monaghan & Ozanne, 2018) and exposure to stressors (Chatelain et al., 2020). Telomere length may reflect somatic integrity, as short telomeres can induce cellular senescence (Blackburn, 2000; Hemann et al., 2001). Telomere shortening also may mirror damage to coding regions of DNA, to the degree that both experience simultaneous assault by oxidative damage (Kawanishi & Oikawa, 2004; von Zglinicki, 2002). Consequently, telomeres are associated with survival among individuals (Frédéric Angelier et al., 2019; Eastwood et al., 2019; Wood & Young, 2019), a pattern shown across vertebrate species, albeit to varying degrees (Wilbourn et al., 2018). One resolution to these mixed results comes from data suggesting that an individual’s *change* in telomere length better predicts survival than telomere length *itself* (Boonekamp et al., 2014; Wood & Young, 2019). These observations challenge the notion that telomeres are causally linked to performance and suggest a vital role for telomere regulatory mechanisms in shaping life history strategies.

At the heart of this issue lies a complex set of processes mediating telomere length. Glucocorticoids, which are linked with survival in many vertebrates (Schoenle et al., 2020), may induce telomere loss via increases in oxidative damage (Kawanishi & Oikawa, 2004; von Zglinicki, 2002) and downregulation of antioxidant defenses (Angelier et al., 2018; Costantini et al., 2011), which buffer telomeres from loss (Badas et al., 2015; Kim & Velando, 2015; Pineda-Pampliega et al., 2020). However, evidence that glucocorticoids increase oxidative damage is mixed (Lendvai et al., 2014; Vagasi et al., 2018; Vitousek et al., 2018). Glucocorticoids also may accelerate telomere loss (reviewed in Angelier et al., 2018), but evidence that this effect is mediated by oxidative stress is stronger *in vitro* than *in vivo* (Boonekamp et al., 2017; Reichert & Stier, 2017), and alternative mechanisms linking glucocorticoids and telomeres are emerging (e.g. mitochondrial metabolism; Casagrande et al., 2020). These data collectively warrant a renewed focus on processes that more directly control telomere length.

Telomerase – and the shelterin proteins that chaperone its effects – orchestrate changes to telomere length. Telomerase is an enzyme that elongates telomeres (Blackburn et al., 1989), and when upregulated, can have a positive effect on somatic health (reviewed in Criscuolo et al., 2018). High telomerase activity drives variation in growth (de Jesus et al., 2011, 2012) and mitochondrial function (Ahmed et al., 2008), but there are potential costs of high telomerase, if it prevents death of unhealthy cells (e.g. tumor growth, Granger et al., 2002; Greider, 1998). Telomerase activity is also stress-responsive (Beery et al., 2012; Choi et al., 2008; Zietzer et al., 2017), and shelterin proteins play an important regulatory role in telomerase activity (de Lange, 2018). Among these is the TPP1-POT1 (‘protection of telomeres 1’) sub-complex, which *physically* interacts with telomerase at the 3’ end, opening and closing the telomere to telomerase activity (Hwang et al., 2012; Wang et al., 2007). While TPP1 recruits telomerase to the telomere (de Lange, 2018), POT1 sterically blocks telomerase from access (Gu et al., 2017; Laprade et al., 2020). Dysregulation of POT1 is therefore common among cancers (reviewed in Wu et al., 2020), contributing to inappropriate cellular immortality. Although telomere elongation in the context of cancer is clearly maladaptive for the organism, comparable processes in healthy tissues may contribute to adaptation by positioning the telomere for repair.

Here, we test the hypothesis that telomere regulatory mechanisms respond to stress and predict performance in wild animals, evaluated with a focus on POT1. Specifically, we ask whether POT1 gene expression is associated with three markers of individual quality in breeding female tree swallows (*Tachycineta bicolor*), including first egg date, body mass, and wing length (Winkler et al., 2020). Next, we experimentally disrupted relationships between POT1 and performance by exposing breeding females to an ecologically relevant stressor (lipopolysaccharide injection), which leads to ‘sickness’ in mothers, and 24h of food restriction in their offspring. We assessed effects of this stressor on both mothers and offspring, including their ability to recover from stress. We measured chick growth and key aspects of telomere biology in chicks (i.e. change in relative telomere length, POT1 gene expression), as well as other traits that have been linked to telomere dynamics in previous work (i.e. glucocorticoids, antioxidants). Because decreases in POT1 should poise the telomere for repair, we predicted that lower POT1 gene expression would be associated with better performance. As we elaborate below, our results introduce POT1 as an ecologically important gene at the intersection of telomere and eco-evolutionary biology.

## 2. METHODS

### 2.1 Study System

This study was approved by Indiana University IACUC #15-004 and all relevant local, state, and federal regulations. We conducted this experiment in spring 2017 in a nest box population we monitor annually in Monroe County, Indiana, USA (39.1851° N, 86.4997° W). This population contains ~270 nest boxes within 15km of one another, including 105 tree swallow nests in 2017 (average brood size = 4.4 ± 0.1). Chicks disperse among these boxes annually, indicating they represent a connected population. We checked boxes regularly to determine lay dates, clutch size, and hatch dates.

We captured adults by hand or nest box trap (Stutchbury & Robertson, 1986) and banded them with a numbered USGS aluminum band on one leg and a colored passive integrated transponder (PIT) tag on the other. Most females were banded during incubation, including 41 of 43 used in this study; n=2 were banded upon treatment. Males were opportunistically banded throughout the breeding season, including 18 of 43 males paired with our focal females. We also recorded data on mass and wing length. Adult sex was confirmed via brood patch or cloacal protuberance.

### 2.2 Experimental Injection of Chick-Rearing Females

We began our experiment on day 5.2 ± 0.1 of the nestling period (hatch day = day 1, range = 3-7 days). We captured females while they provisioned, either by blocking the entrance hole while the female was inside or by setting a nest box trap. We weighed each female and collected ≤50μL blood for RNA. Following Palacios et al. (2011), we administered a subcutaneous injection of either saline or lipopolysaccharide (LPS) saline-oil emulsion in the right dorsal apterium (saline: n=22, LPS: n=21; see **SI materials**). LPS is a non-replicating piece of bacterial cell wall commonly used to temporarily trigger an immune response and induce ‘sickness’ behaviors, e.g. lethargy, weight loss, reduced parental care (Dantzer et al., 2008; Palacios et al., 2011). We recaptured 37 of the 43 females 24-48h later (n=29 within 24h, saline: n=16, LPS: n=13), at which time we weighed them and collected another ≤50μL blood for RNA. Percent body mass change in the 24h following treatment was calculated relative to starting mass: ((mass _post_ – mass _pre_)/mass _pre_) × 100. We could not recapture 6 females, including 2 who abandoned this breeding attempt.

### 2.3 Visitation Rates

Past work shows that parental visitation rate is a reliable proxy of provisioning (McCarty, 2002), and we used radio-frequency identification (RFID) boards to measure visitation rate. All nest boxes in the study were equipped with RFID readers, which recorded a unique tag ID and time stamp every time a PIT tag passed through the antennae at the box entrance (Bonter & Bridge, 2011; Lendvai et al., 2011). We determined the number of visits by filtering out continuous reads occurring within 3 sec of another read of the same individual, often generated when a bird is perched at the nest entrance. To account for entrances and exits, we halved the number of remaining reads. Previous work suggests that LPS-induced sickness can last 48h, but peak effects occur around 3-6h post-injection (Dantzer et al., 2008). Therefore, we quantified visitation rate as the average hourly number of visits occurring 3-6h post-injection. Baseline visitation rates were taken the day prior during the same 3h window (n=35; n=8 excluded due to equipment failure). We also used 30-min behavioral observations to confirm that RFID and observed visitation rates were correlated (F_1,53_=6.14, p=0.0168, n=55, r=0.903; see **SI materials**).

### 2.4 Measuring Phenotypic Effects on Chicks

We assessed how maternal treatment affected chicks in the subsequent week and limited our analyses to those nests treated during peak chick growth, which occurs at 5 to 6-days old (Wolf et al., in press). We therefore excluded 3 nests with 4 or 7-day old chicks and 3 additional failed nests, which left a total of 37 nests (saline: n=20, LPS: n=17). While mothers were being injected, we measured nestling mass to the nearest 0.1g using an electronic scale, collected a chick blood sample from the metatarsal vein (30-50μL), and gave each chick a unique nail trimming for later identification (n=167 total chicks). The following day, we reweighed all chicks to calculate percent body mass change in the 24h following maternal treatment: ((mass _post_ – mass _pre_)/mass _pre_) × 100.

We visited nestlings a third time when they were 12-days old. At this age, chicks are approaching asymptotic adult-like mass (Wolf et al., in press), they exhibit adult-like corticosterone secretion (Wada et al., 2007), and they are not likely to prematurely fledge. During sample collection, we left one chick in the nest to minimize disturbance to parents. For remaining chicks, we sought to collect blood from the alar vein within 3 min of disturbance (latency: 2:08 ± 0:44 min) to measure baseline circulating corticosterone (hereafter CORT), and again at 30 min (31:14 ± 1:00 min) to measure handling-induced elevated CORT. For the subset of nests whose mothers were treated when chicks were 5-days old, we opportunistically collected an additional blood sample within 5 min of disturbance (1:24 ± 1:12 min) for gene expression analyses (Herdegen & Leah, 1998). Due to logistical constraints, we did not obtain each blood sample from every chick, as elaborated below. In total, we collected ≤200μL blood from each chick, below the maximum suggested volume (Gaunt et al., 1997), based on an average 12-day old body mass of 19.9±0.2g. We banded all chicks with one numbered USGS band. Blood was stored on ice (for hormones and DNA) or dry ice (for RNA). Later the same day, we centrifuged hormone samples, reserved plasma, and stored at −20°C. Whole blood and red blood cells were stored at −80°C.

At ~21 days post-hatch, we inspected all nests for signs of fledging (e.g., flattened nest, feces accumulation) or failure (e.g., remains, disturbed nest). We identified any remaining (dead) chicks based on leg bands, and assumed other chicks successfully fledged if the nest showed no signs of failure, following best practices in avian field biology (Martin & Geupel, 1993; McCarty, 2001). During the following two breeding seasons (2018-2019), we captured breeding birds to estimate recruitment of chicks into the breeding population, as has been done in previous work in tree swallows (Lombardo et al., 2020; Shutler et al., 2006). We devoted substantial monitoring effort from March to July each year to locate and capture returning chicks, which can be easily distinguished by their single aluminum band.

### 2.5 Quantifying Plasma Corticosterone

We quantified plasma CORT using an enzyme immunoassay kit (Cayman #501320; assay sensitivity = 30 pg/mL), which we previously validated in tree swallows (Virgin & Rosvall, 2018). We used 107 chicks from 34 nests (saline: n=20, LPS: n=14) for which we obtained sufficient plasma (>10μL) for both baseline and 30 min sampling points. We combined each 10μL plasma with 200μL dH_2_0, vortexed, and performed 3 rounds of ether extractions. We dried extracts with N_2_ and reconstituted with 600μL assay buffer. While we did not correct for extraction efficiency, recoveries are likely high because our extraction protocols have previously shown >90% efficiencies (George & Rosvall, 2018). Each plate included the following in duplicate: 8-point standard curve, blank, maximum binding, non-specific binding, total activity, 3 plasma pools (for intra-and inter-plate variation), and 33 samples. We ran samples across 10 plates, balanced by date, treatment, mass, and brood size. We read absorbance at 412nm using an Epoch spectrophotometer (BioTek, Winooski, VT, USA) and interpolated CORT levels using Gen5 software (v.2.09.2, BioTek). Inter-plate coefficient of variation (CV) was a 12.9% and intra-plate CV was 5.1 ± 1.8%.

### 2.6 Quantifying Gene Expression

Our primary goal was to measure gene expression of POT1. We also generated a molecular measurement of antioxidant capacity (as in Sridhar et al., 2014; Yarru et al., 2009) – instead of measuring total antioxidant capacity in plasma using the OXY-ADSORBENT test (as in Beaulieu et al., 2011) – because plasma was depleted in CORT analyses. Specifically, we quantified the gene expression of glutathione peroxidase (GPX), peroxiredoxin-1 (PRDX-1), and superoxide dismutase (SOD). The products of these genes influence multiple measures of antioxidant function (Pisoschi & Pop, 2015), suggesting they are likely to be a generalized measure of antioxidant capacity. Gene expression values for all three antioxidants were positively correlated (R^2^ > 0.62), so we used a principal components analysis to condense these data. PC1 (Eigenvalue = 1.58) negatively loaded for all three antioxidants (SOD: −0.56, GPX: −0.61, PRDX1: −0.57) and accounted for 83% of the total variance. PC1 was multiplied by −1 so that positive values indicate higher gene expression. All gene expression data was log2-transformed for analyses.

We extracted RNA from whole blood using a phenol-chloroform-based Trizol method (Invitrogen, Carlsbad, CA) using PhaseLock tubes (5PRIME, #2302830). We synthesized cDNA using 1μg RNA and Superscript III reverse transcriptase (Invitrogen), treated with DNAase (Promega, Madison, WI) and RNase inhibitor (RNAsin N2111, Promega). cDNA was stored at −20°C. For each gene of interest, we used the 2^−ΔΔCT^ method of quantitative PCR, in which expression is normalized against a reference gene and relative to a calibrator sample run on each plate. We used PPIA (peptidylprolyl isomerase A) as a reference gene, as it is highly expressed in blood and reliable in birds (Zinzow-Kramer et al., 2014). All primer sequences were developed from the tree swallow transcriptome (accession #GSE126210; Bentz et al., 2019), and further details are reported in **Table S1**. Samples were run alongside no template controls, using PerfeCta SYBR Green FastMix with low ROX (Quanta Biosciences, Gaithersburg MD) on 384-well plates using an ABI Quantstudio 5 machine (Thermo Fisher Scientific, Foster City, CA) with Quantstudio Design & Analysis software (v1.4.3, Thermo Fisher Scientific). Each well included 3μL of cDNA diluted 1:50 (or 3μL water, for NTCs) and primers diluted to 0.3μM in a total volume of 10μL. All reactions use the following thermal profile: 10 min at 95°, followed by 40 cycles of 30 s at 95°, 1 min at 60°, and 30 s at 70°, with a final dissociation phase (1 min at 95°, 30 s at 55°, and 30 s at 95°) that confirmed single-product specificity for all samples. All samples fell within the bounds of the standard curve and the reaction efficiencies were always within 100 ± 15%. A pool reference sample present on all plates was used to calculate intra- and inter-plate variation. Samples were run in triplicate, and the mean values were used to calculate the relative quantity for each sample using the following formula: 2^−ΔΔCt^, where ΔΔCt = (Ct ^GOI^ – Ct ^PPIA^)_reference_ – (Ct ^GOI^ – Ct ^PPIA^)_sample_. Mean intra and inter-plate variation of the C_t_ values were 0.359% and 1.18% for PPIA, 0.156% and 0.397% for SOD, 0.315% and 0.636% for PRDX-1, 0.681% and 0.437% for GPX, and 0.558% and 2.34% for POT1. In total, we quantified gene expression for n = 77 nestlings (saline: n=48, LPS: n=29), taken from 26 nests (saline: n=15, LPS: n=11).

### 2.7 DNA Extraction and Molecular sexing of chicks

We used the automated Maxwell® RSC Instrument (Promega, Madison, WI) and Whole Blood DNA Kit (#AS1520) to extract DNA from ≤25μL red blood cells. We determined the sex of all chicks following established methods (Çakmak et al., 2017). Males exhibited a single band at ~250bp and females exhibited a double band at ~250 and 275bp (see **SI Material**).

### 2.8 Telomere Measurement

We quantified relative telomere length using qPCR, adapted from (Cawthon, 2009; Criscuolo et al., 2009). Relative telomere length was measured as the ratio (T/S) of telomere repeat copy number (T) to a single gene copy number (S), relative to a pooled reference sample present on all plates. We amplified our single copy gene, glyceraldehyde-3-phosphate dehydrogenase (GAPDH) and telomeres using primers telg/telc (see **Table S1**). We conducted qPCR on 384-well plates (ABI Quantstudio 5, Foster City, CA). For each sample, we ran GAPDH and telomere reactions on the same plate. Prior to plating, we diluted DNA samples to 3.33ng/μL using ultra-pure water. Each reaction had a total volume of 10μL containing 5μL PerfeCTA SYBR Green SuperMix Low ROX (Quanta Biosciences, Gaithersburg, MD, USA), 200nM each GAPDH-F/GAPDH-R or 200nM each telc/telg, and 3μL DNA extract (10ng total). qPCR reaction conditions were: 10 min at 95°C, followed by 30 cycles of 10 s at 95°C, 1 min at 62°C, and 30 s at 72°C, followed by 1 min at 95°C, 30 s at 55°C, and 30 s at 95°C. In both reactions, the number of PCR cycles necessary to accumulate sufficient fluorescent signal to cross a threshold (C_t_) was measured and individuals with relatively longer telomeres were characterized by shorter reaction times. All samples fell within the bounds of the standard curve and the reaction efficiencies were always within 100 ± 15% (GAPDH: 98.7 ± 2.8; telomeres: 107.6 ± 8.2). A tree swallow pool reference sample present on all plates was used to calculate intra- and inter-plate variation. Samples were run in triplicate, and mean values were used to calculate T/S ratios for each sample using the formula: 2^−ΔΔCt^, where ΔΔC_t_ = (C_t_ ^telomere^ – C_t_ ^GAPDH^)_reference_ – (C_t_ ^telomere^ – C_t_ ^GAPDH^) _reference_. From the original 167 chicks in the study, 161 survived to 12-days old. From those 161 chicks, 147 have telomere measurements from both pre-treatment and 12-days old (saline: n=87, LPS: n=60). From the 14 missing chicks, 9 were from 2 LPS nests inadvertently sampled at 11 or 13-days old, and 5 chicks had poor replicates for one or both samples and were excluded. Telomere attrition was corrected for regression to the mean (Verhulst et al., 2013), where more negative values indicate greater telomere loss.

Intraplate and interplate repeatabilities were calculated using the R package ‘rptr’ (Stoffel et al., 2017). Intraplate repeatability (estimated via intraclass correlation coefficient) was 0.96 ± 0.004 (95% CI = 0.95, 0.97) for GAPDH C_t_ values, 0.88 ± 0.011 (95% CI = 0.86, 0.90) for telomere C_t_ values, and 0.76 (95% CI: 0.72, 0.80) for 2^−ΔΔCt^ measurements. Interplate repeatability of 2^−ΔCt^ for reference samples was 0.84 ± 0.13 (95% CI = 0.46, 0.94). Because pre- and post-treatment pairs of samples were run on the same plate for each individual, and treatment was balanced across plates, plate effects on 2^−ΔΔCt^ values should be minimal; however, we accounted for plate ID in telomere analyses to control for potential plate effects.

Next, we conducted a sensitivity analysis to assess the degree of measurement error in our telomere variable. We tested whether the random effect estimate of individual ID explained more variance in relative telomere length among technical replicates, i.e. triplicates next to each other on a plate, than variance between biological replicates, i.e. from pre-treatment to 12-days old (similar to van Lieshout et al., 2019, see SI XXX for details). After accounting for plate effects using MCMCglmm (Hadfield, 2010), the random effect estimate for individual ID explained more variance in relative telomere length among technical replicates (0.077; 95% CI = 0.057, 0.098) than among biological replicates (0.0044; 95% CI = 0.0001, 0.01), meaning that noise in telomere measurements between technical replicates was much lower than biologically-relevant changes occurring within an individual during the study. We also parsed the data based on whether an individual’s telomere length change was positive or negative to separately analyze technical and biological variance within these groups. None of these analyses show overlap of 95% CI for random effect estimates between technical and biological replicates (**Figure S1**), indicating that our measure is highly precise.

### 2.9 Statistical Analyses

All statistical analyses were performed in R (version 3.5.3, R Core Team, 2019). We used an information-theoretic approach to evaluate support for competing candidate models predicting each variable of interest. For each dependent variable, we used *dredge* (Barton, 2019) to create model sets from the global model (detailed below), in which all models for a given response variable included the same subset of data. For global models, we assessed multicollinearity and removed redundant variables with variable inflation factors ≥ 5 (Fox & Weisberg, 2011). We used Akaike Information Criterion (AIC_c_ – to correct for sample size) for model comparisons (Burnham & Anderson, 2002), and we present ΔAIC (AIC_i_ – AIC_best model_) and AIC weights (weight of evidence for model) for highly supported models with ΔAIC ≤ 2 compared to the top model (K. P. Burnham et al., 2011). All models with ΔAIC ≤ 2 are equally fit, so when this occurred, we report the most parsimonious model (K. Burnham & Anderson, 2002). To estimate how well models fit our data, we calculated pseudo-R^2^ using the MuMIn package (Barton, 2019), which considers variance explained by either fixed (R^2^_marginal_) or both fixed and random effects (R^2^_conditional_). Variable significance of top or most parsimonious models was assessed using restricted maximum likelihood. We visually inspected model residuals for normality and homoscedasticity.

To ask whether POT1 gene expression or relative telomere length in the blood has more overall support in predicting traits of interest, we directly compared models including POT1 gene expression vs. relative telomere length. If multiple models were supported for each trait, we calculated variable importance of each predictor variable, or the sum of AIC_c_ weights for models containing that variable, where predictor variables with a value of 1 indicate greatest importance.

#### 2.9.1 Does POT1 or relative telomere length better predict female quality?

We tested the hypothesis that POT1 gene expression and relative telomere length predict metrics of quality in breeding females, including first egg date, body mass, and wing length. For each trait, we ran a series of models which assumed a gaussian distribution and contained all combinations of POT1 gene expression, relative telomere length, first egg date (except when first egg date was the response variable), chick age at the time of the female’s capture, and brood size. Analyses utilized 37 females for which we had all predictor variables.

#### 2.9.2 Does POT1 or relative telomere length better predict female responses to stress?

We next tested the hypothesis that POT1 gene expression and relative telomere length predict a female’s response to LPS injection, specifically her own change in body mass and visitation rate during sickness. For each trait, we ran a series of models that assumed a gaussian distribution and contained all combinations of POT1 gene expression, relative telomere length, treatment, first egg date, chick age, and brood size. Treatment was included in all candidate models. Analyses of mass change included the n=24 females for whom we have all predictor variables and recaptured within 24h of injections. Analyses of visitation rate included n=31 females for whom we have all predictor variables.

#### 2.9.3 How does maternal stress influence chick phenotypes?

##### Evaluating correlations among traits

We evaluated correlations between phenotypic qualities in chicks (i.e., growth, change in relative telomere length, POT1 and PC1 for antioxidant gene expression, baseline and handling-induced CORT). We visualized relationships within each treatment using the corrplot package (Wei et al., 2017) and computed Spearman’s *r* and p-values, adjusted for false discovery (Benjamini & Hochberg, 1995).

##### Treatment effects on chick phenotypes

We next tested how maternal LPS treatment affected chick phenotypes, including growth during the 24hr following maternal injection, telomere attrition from pre-treatment to 12-days old, POT1 gene expression, PC1 for antioxidant gene expression, baseline CORT, and handling-induced CORT. Because traits were largely uncorrelated (**Fig S4**), we analyzed each trait separately. Two outliers were detected in the chick growth dataset (Grubb’s test, p<0.05) and were removed. To meet model assumptions, baseline and handling-induced CORT were log-transformed. For each trait, we ran a series of linear mixed-effects models using the nlme package (Pinheiro et al., 2019) that assumed a gaussian distribution and included main effects of treatment, sex, treatment x sex interaction, brood size, hatch date, and a random effect of nest. For CORT models, we also included a fixed effect of time of day, and for change in relative telomere length models, we included qPCR plate as a fixed effect. Because our primary question asks how treatment influences these traits, treatment was included in all candidate models, and model comparisons allowed us to assess which covariates to include in final analyses. We report Satterthwaite-adjusted degrees of freedom. Due to logistical considerations described above, final sample sizes per global model varied by response variable: n=159 chick growth; n=147 change in relative telomere length; n=77 POT1 gene expression; n=77 PC1 for antioxidant gene expression; n=107 baseline CORT; n=107 30-min CORT.

#### 2.9.4 How do chick phenotypes relate to survival?

To test how chick phenotypes relate to survival to fledging or recruitment the following year, we ran a series of generalized linear mixed models which assumed a binomial distribution and included treatment, sex, change in relative telomere length, POT1 and PC1 for antioxidant gene expression, 12-day old body mass, baseline CORT, and handling-induced CORT, with nest box as a random effect. As above, CORT data were log-transformed. The analysis on survival to fledging included 76 chicks from 26 broods for which we had all physiological metrics taken at 12-days old. Of these 76 chicks, only 11 failed to fledge. The analysis on recruitment included 65 fledged chicks from 26 broods for which we had all physiological metrics. Of these 65 chicks, only 5 recruited into the breeding population the following year, typical of the 5-10% return rate in this species (Winkler et al., 2020). PC1 for antioxidant gene expression was removed from the global model for recruitment to avoid multicollinearity with POT1 (variable inflation = 9.21).

## 3. RESULTS

### 3.1 POT1 gene expression and female quality

POT1 gene expression largely outperformed relative telomere length as a predictor of variation in female quality. Top-ranked models for first egg date and body mass both contained POT1 gene expression but not relative telomere length (**Table 1A**), and POT1 gene expression had higher variable importance than relative telomere length when summed across all candidate models (**Fig 2**, **Fig S2**). The best supported model predicting first egg date contained POT1 gene expression: earlier breeding females exhibited lower POT1 gene expression (F_1,35_=4.80, p=0.035; **Table 1A; Fig 1A**). The top model for body mass indicated significant main effects of POT1 gene expression and chick age, where body mass was heavier in females with lower POT1 gene expression (F_1,34_=5.25, p=0.028) and those with younger chicks (F_1,34_=4.65, p=0.038; **Table 1A; Fig 1B**). The top-ranked model predicting wing length included first egg date, but the null model was most parsimonious (**Table 1A; Fig 1C**).

**Table 1.** The top models (ΔAIC_c_ ≤ 2) assessing the role of POT1 gene expression (POT1) and relative telomere length (RTL) in predicting (A) metrics of female quality and (B) maternal responses to injection. Global models are summarized in Section 2.9. K = # of parameters (including intercept), w_i_ = model weight, FED = first egg date.

| Variable | Model | K | AICc | $\Delta AIC_c$ | $w_i$ | $R^2_m$ |
| --- | --- | --- | --- | --- | --- | --- |
| **(A) Metrics of female quality** |  |  |  |  |  |  |
| **First Egg Date**<br>(n=37) | POT1 | 2 | 239.98 | 0 | 0.252 | 0.118 |
|  | POT1 + Brood Size | 3 | 241.4 | 1.42 | 0.124 | 0.14 |
|  | POT1 + Chick Age | 3 | 241.51 | 1.53 | 0.117 | 0.137 |
|  | POT1 + RTL | 3 | 241.58 | 1.6 | 0.113 | 0.135 |
| **Body Mass**<br>(n=37) | POT1 + Chick Age | 3 | 121.25 | 0 | 0.281 | 0.216 |
|  | POT1 + Chick Age + FED | 4 | 123.06 | 1.81 | 0.114 | 0.228 |
|  | POT1 + Chick Age + RTL | 4 | 123.11 | 1.86 | 0.111 | 0.227 |
| **Wing Length**<br>(n=37) | FED | 2 | 178.79 | 0 | 0.135 | 0.074 |
|  | Intercept only (null) | 1 | 179.33 | 0.54 | 0.103 | 0 |
|  | RTL | 3 | 179.6 | 0.81 | 0.09 | 0.112 |
|  | POT1 + FED | 3 | 179.79 | 1 | 0.082 | 0.107 |
|  | RTL | 2 | 179.82 | 1.03 | 0.081 | 0.048 |
|  | Brood + FED | 3 | 180.74 | 1.95 | 0.051 | 0.085 |
| **(B) Maternal responses to injection** |  |  |  |  |  |  |
| $\Delta$ **Body Mass**<br>(n=24) | Treatment + RTL | 3 | 136.25 | 0 | 0.176 | 0.15 |
|  | Treatment | 2 | 136.36 | 0.11 | 0.167 | 0.048 |
|  | Treatment + FED | 3 | 137.45 | 1.2 | 0.097 | 0.11 |
|  | Treatment + POT1 + RTL | 4 | 137.57 | 1.32 | 0.091 | 0.203 |
|  | Treatment + FED + RTL | 4 | 137.77 | 1.52 | 0.082 | 0.197 |
| $\Delta$ **Visitation Rate**<br>(n=31) | Treatment*POT1 | 4 | 163.2 | 0 | 0.365 | 0.36 |
|  | Treatment + POT1 | 3 | 164.85 | 1.65 | 0.16 | 0.274 |

**Fig 1.**
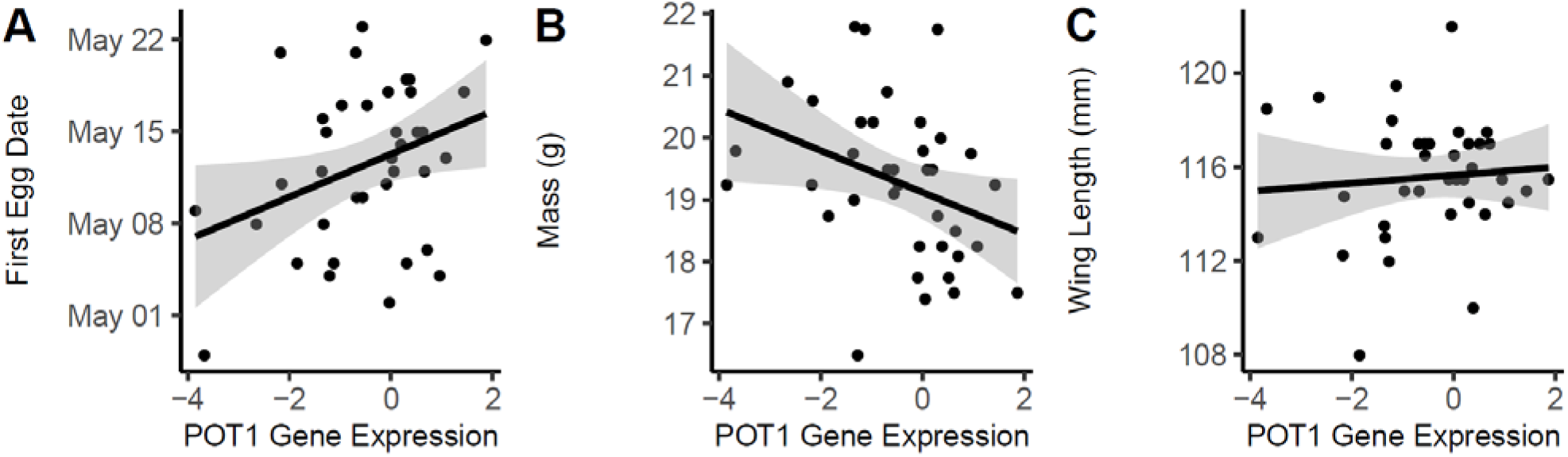
POT1 gene expression predicts metrics of female quality, including first egg date (A), body mass (B), but not wing length (C). Shading indicates 95% confidence interval from model output.

### 3.2 POT1 gene expression and female responses to stress

Female POT1 gene expression was unaffected by treatment (F_1,24_=0.086, p=0.77), and pre-injection and post-injection mRNA abundances were positively correlated (ρ=0.76, p<0.0001), suggesting consistency in POT1 gene expression in our adult study subjects.

For models predicting treatment-induced changes in visitation rate, POT1 gene expression had more overall support than relative telomere length (**Fig 2, Fig 3B**, **Fig S3B**). The top-ranked model predicting changes to visitation rate indicated a treatment x POT1 interaction showing that among LPS-injected females, those with the lowest POT1 gene expression best maintained high visitation rates (**Fig 3B**). However, the most parsimonious model included only treatment and POT gene expression: LPS-injected females significantly decreased visitation rates during the peak of sickness (saline = −0.67 ± 0.59 visits/hour; LPS = −3.74 ± 1.08 visits/hour; F_1,28_=7.44, p=0.011, **Table 1B, Fig 3B**), and females with the lowest POT1 gene expression exhibited marginally higher visitation rates (F_1,28_=3.89, p=0.059).

**Fig 2.**
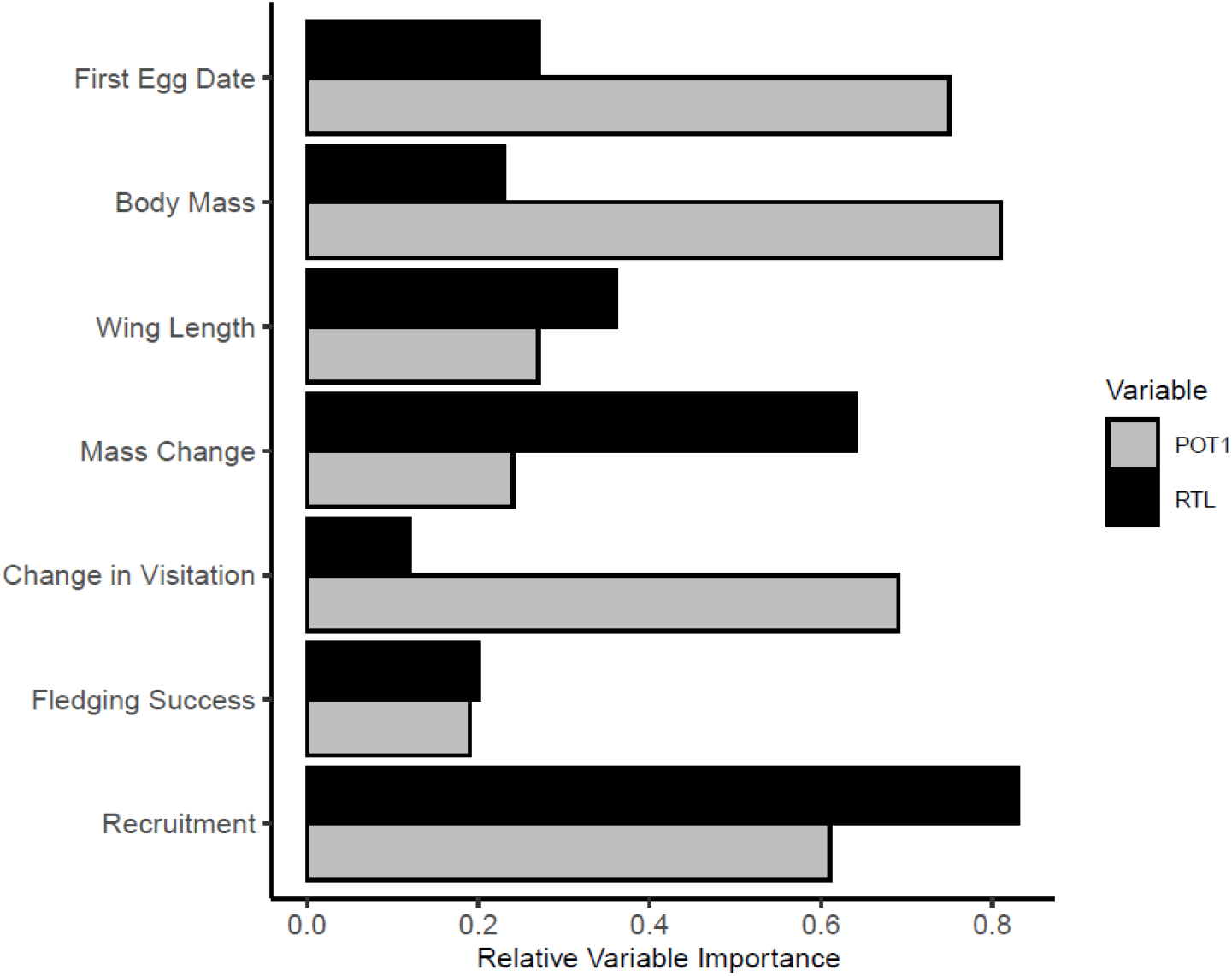
Comparing the importance of relative telomere length (RTL, black) and POT1 gene expression (gray) in predicting variation in fitness-related traits. Variable importance is the sum of Akaike weights across all models in which the variable occurred.

**Fig 3.**
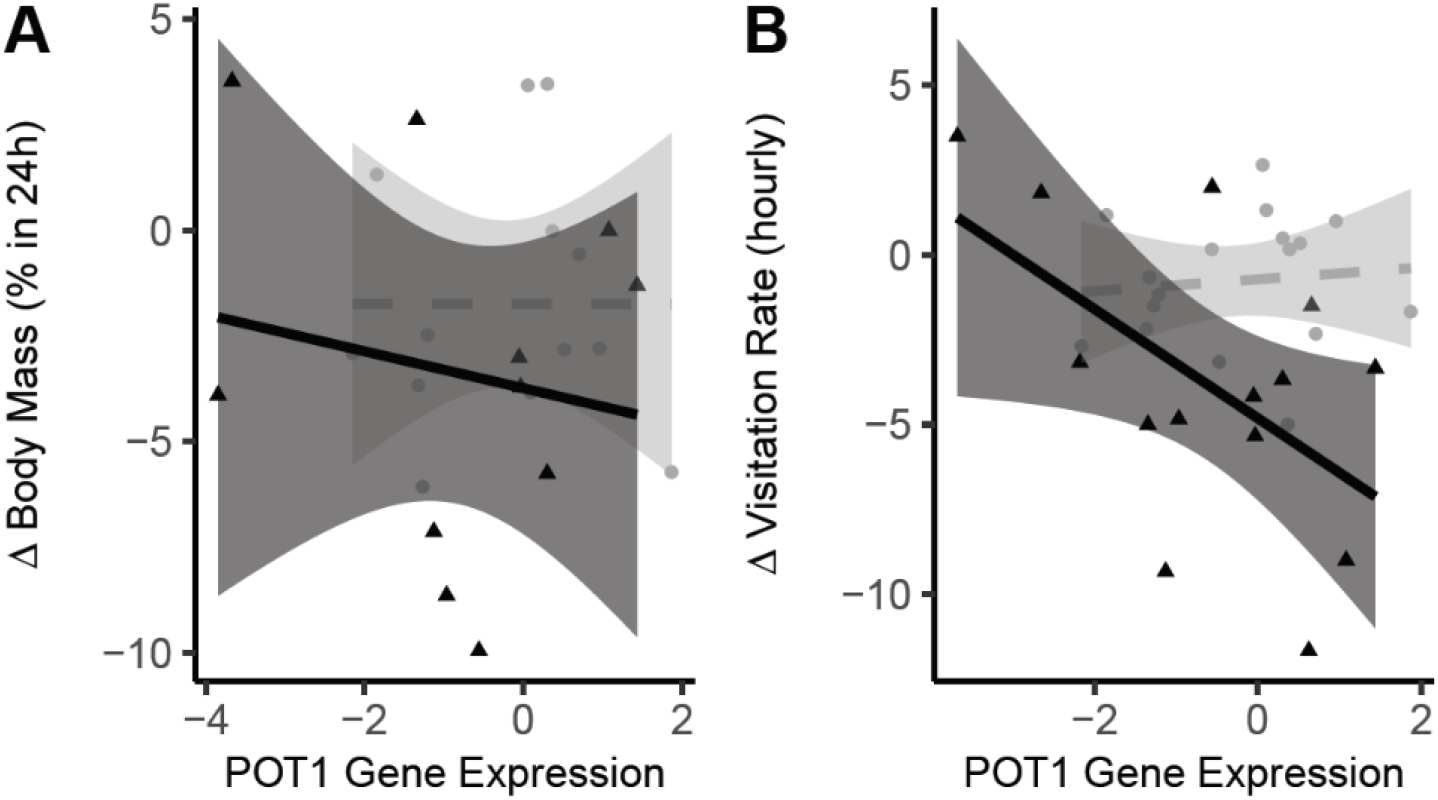
Female POT1 gene expression and responses to stress following injection of breeding mothers with saline (gray, circles, dashed line) or LPS (black, triangles, solid line): A) female Δ body mass within 24h of injection or B) Δ visitation rate during the peak of sickness.

For models predicting a female’s treatment-induced change in body mass, relative telomere length had more overall support (higher variable importance) than POT1 gene expression. While the top-ranked model contained treatment and relative telomere length, telomere length was not a significant predictor of mass change (F_1,21_=2.81, p=0.11), and the most parsimonious model included treatment alone (**Table 1B; Fig 3A**, **Fig S3A**). There was no significant treatment effect on body mass (F_1,22_=1.16, p=0.29), though LPS-treated females varied from a loss of 9.9% to a gain of 3.5% body mass (**Fig 3A**).

### 3.3 Effects of maternal injection on chick phenotypes

#### Relationships among phenotypes

Within saline and LPS chicks, we only found one significant association between telomere-related mechanisms: there was a positive correlation between POT1 gene expression and PC1 for antioxidant gene expression (**Fig S4A**, **Fig S4B**).

#### Differences in chick phenotypes in response to maternal injections

While the top-ranked model predicting chick growth included main effects of treatment, sex (and their interaction), and brood size, the most parsimonious model predicting chick growth contained only treatment, sex, and brood size (**Table S2**). Chicks of LPS-females grew significantly less in the 24h following maternal injection (F_1,32_=11.98, p=0.0015, **Fig 4A**, **Table S2**), with no main effects of sex (F_1,122_=0.99, p=0.32) or brood size (F_1,33_=2.02, p=0.16).

**Fig 4.**
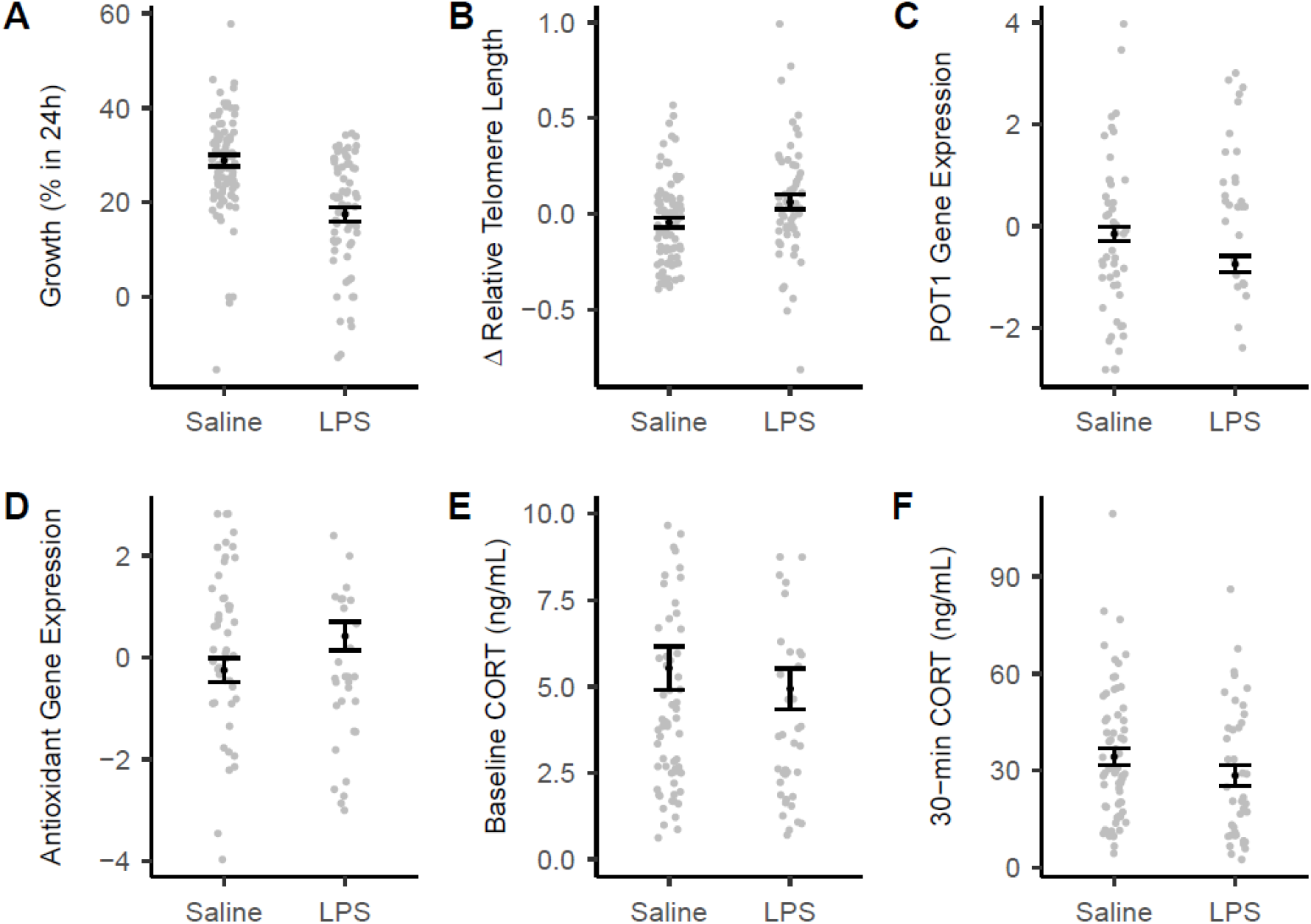
The effects of maternal treatment on chick phenotypes. A) Growth % during the 24h following maternal injection, B) change in relative telomere length, C) POT1 gene expression, D) antioxidant gene expression (i.e., PC1 for superoxide dismutase, glutathione peroxidase, and peroxiredoxin-1), E) baseline CORT, and F) handling-induced CORT. Gene expression data are on a log-2 scale. Global models included treatment, sex, treatment x sex interaction, hatch date, and brood size, with a random effect of nest; see ***SI Table S2***. Errors bars are mean ± SE.

The top-ranked model predicting change in relative telomere length contained treatment and qPCR plate (**Fig 4B**, **Table S2**), where chicks of LPS-injected females exhibited significantly less telomere shortening relative to controls (F_1,32_=4.19, p=0.049, plate effect: F_1,32_=0.36, p=0.55), which cannot be explained by measurement error alone (see **SI material**).

The top-ranked and most parsimonious model predicting POT1 gene expression included treatment and sex: POT1 gene expression was significantly lower in chicks of LPS-injected females (F_1,24_=6.06, p=0.02, **Fig 4C**, **Table S2**) and in female chicks (F_1,50_=6.52, p=0.014).

The top-ranked model predicting PC1 for antioxidant gene expression indicated significantly lower POT1 gene expression in female chicks regardless of treatment, but the most parsimonious model included treatment as the only main effect: chicks of saline and LPS-injected females did not significantly differ in antioxidant gene expression (F_1,24_=2.13, p=0.16, **Fig 4D**, **Table S2**).

Although the top-ranked model predicting baseline CORT included treatment and time of day, the most parsimonious model showed that baseline CORT did not significantly differ by treatment (F_1,32_=0.39, p=0.54, **Fig 4E**, **Table S2**). The top-ranked model predicting handling-induced CORT showed no significant effects of treatment (F_31_=2.53, p=0.12, **Fig 4F**, **Table S2**) or time of day (F_1,31_=2.14, p=0.15).

### 3.4 Model comparisons linking chick phenotypes to survival

The highest-ranking and most parsimonious model predicting chick survival to fledging included mass and handling-induced CORT at 12-days old (**Table 2)**. Compared to chicks that did not survive, fledged chicks had a heavier body mass (z=2.81, p=0.005) and marginally higher handling-induced CORT secretion (z=1.88, p=0.059). These same traits had high importance across all candidate models predicting fledging success, while variable importance of change in relative telomere length and POT1 gene expression were low (**Fig 2**).

**Table 2.** The top models (ΔAIC_c_ ≤ 2) assessing the role of POT1 gene expression and relative telomere length (ΔRTL) in predicting chick fledging and recruitment into the breeding population. The global model included treatment, sex, ΔRTL, POT1 and PC1 antioxidant gene expression, 12-day old body mass (Mass), and log-transformed baseline CORT (T0 CORT) and handling-induced CORT (T30 CORT). All models include the intercept and random effect of nest. K = # of parameters (including intercept), w_i_ = model weight, R^2^_m_ = variance explained by fixed effect, R^2^_c_ = variance explained by fixed and random effects.

| Variable | Model | K | AICc | $\Delta AIC_c$ | $w_i$ | $R^2_m$ | $R^2_c$ |
| --- | --- | --- | --- | --- | --- | --- | --- |
| **Fledging Success**<br>**(n=76)** | Mass + T30 CORT | 3 | 51.17 | 0 | 0.116 | 0.466 | 0.466 |
|  | Mass + T30 CORT + Sex | 4 | 51.45 | 0.28 | 0.1 | 0.515 | 0.635 |
|  | Mass + T30 CORT + T0 CORT + Sex | 5 | 52.62 | 1.45 | 0.056 | 0.522 | 0.522 |
|  | Mass + T30 CORT + T0 CORT | 4 | 53.04 | 1.87 | 0.045 | 0.469 | 0.469 |
| | $\Delta RTL$ + Mass + T30 CORT + Sex | 5 | 53.08 | 1.91 | 0.044 | 0.518 | 0.599 |
| | $\Delta RTL$ + Mass + T30 CORT | 4 | 53.13 | 1.96 | 0.043 | 0.473 | 0.473 |
|  | POT1 + Mass + T30 CORT | 4 | 53.16 | 1.99 | 0.043 | 0.467 | 0.467 |
| **Recruitment**<br>**(n=65)** | $\Delta RTL$ + POT1 | 3 | 34.51 | 0 | 0.172 | 0.475 | 0.541 |
| | $\Delta RTL$ + POT1 + Mass | 4 | 36.11 | 1.6 | 0.077 | 0.516 | 0.516 |
| | $\Delta RTL$ + POT1 + Sex | 4 | 36.11 | 1.6 | 0.077 | 0.544 | 0.544 |
| | $\Delta RTL$ + POT1 + Treatment | 4 | 36.27 | 1.76 | 0.071 | 0.498 | 0.585 |
| | $\Delta RTL$ | 2 | 36.34 | 1.83 | 0.069 | 0.292 | 0.292 |

Recruitment models pointed to a different set of traits. The top-ranked model for recruitment into the breeding population included main effects of both change in relative telomere length and POT1 gene expression, but the most parsimonious model included only change in relative telomere length: regardless of treatment, chicks experiencing less telomere shortening were more likely to recruit into the breeding population the following year (z=2.18, p=0.030; **Table 2**). Across all candidate models predicting recruitment, change in relative telomere length and POT1 gene expression had the highest importance, 0.83 and 0.61, respectively (**Fig 2**).

## 4. DISCUSSION

In the field of ecology and evolution, long telomeres have become nearly synonymous with a variety of fitness-related traits, yet very little work considers the role of telomere regulatory mechanisms in adaptive phenotypic plasticity in nature. To our knowledge, this is the first study to assess how natural variation in gene expression of a shelterin protein predicts performance in a free-living system. Lower levels of POT1 gene expression should facilitate telomeric repair, and we show low mRNA abundance is related to metrics of high quality in adult females. Experimental evidence echoes this view: females with naturally lower POT1 gene expression were most able to maintain parental care during an ecologically relevant stressor that otherwise decreased provisioning and temporarily slowed chick growth. Interestingly, chicks of LPS-injected mothers decreased POT1 gene expression and exhibited telomere elongation in the following week, consistent with the idea that low POT1 permits telomere buffering from stress. We also find some support that these effects are visible to natural selection: changes in telomere length and POT1 gene expression were the strongest predictors of chick recruitment the following year. Together, our results suggest that telomere regulators are stress-responsive and predict performance, oftentimes more so than telomere length itself. As we discuss below, telomere regulatory genes like POT1 are potentially key modulators of variation in life history strategies.

### POT1 gene expression predicts performance in breeding females

Raising young is a predictably challenging life history stage, and we show that adult females with lower POT1 gene expression exhibited higher quality traits, including an earlier start to breeding and a heavier body mass during the chick period, both of which are condition-dependent traits associated with higher reproductive performance in tree swallows (Winkler et al., 2020). Our experimental manipulation also suggests that POT1 predicts a female’s handling of stress: LPS-injected females decreased in provisioning rates, but this effect was weakest for constitutively low-POT1 females, who exhibited visitation rates comparable to those of saline-injected controls. Critically, POT1 gene expression showed higher variable importance than relative telomere length in predicting most of these performance-related traits, and variables better predicted by telomeres – wing length and female mass change – were not significantly related to relative telomere length (see **Fig S2C, S3A**). These results contribute to a growing body of literature linking telomeres with timing of breeding (Bauch et al., 2013; Bauer et al., 2018; Le Vaillant et al., 2015) and body mass (Angelier et al., 2015; Angelier et al., 2019). Our findings therefore extend this work with new perspectives in which telomere *regulation* may track fitness-related traits.

### POT1 gene expression predicts responses to acute stress in chicks

While stress-responsive elements, such as glucocorticoids or antioxidants, may affect telomerase *expression* (Beery et al., 2012; Choi et al., 2008; Zietzer et al., 2017), POT1 should alter telomerase *efficacy* because of competitive binding between telomerase and POT1 at the 3’ telomere end. Consequently, decreased POT1 gene expression in chicks of LPS-treated females may increase telomerase access to the telomere and buffer telomeres from loss. This idea is consistent with the relative telomere elongation seen in stress-exposed chicks, but whether POT1 covaries quantitatively with telomerase activity requires further study, considering that we did not observe a linear relationship between POT1 mRNA abundance and telomere dynamics (**Fig S4**). Biomedical work shows POT1 gene expression to be plastic following stress in rodents and stem cells (Ludlow et al., 2017; Ludlow et al., 2012; Moazzam et al., 2020), but we are not aware of comparable results in a wild animal. Interestingly, POT1 gene expression did not respond to the LPS-induced stressor in adult females, suggesting that plasticity in POT1 is context-dependent or only occurs when telomerase activity is also high, such as the early postnatal period (Haussmann et al., 2007). How we interpret these dynamics in blood calls for a closer look at blood-producing tissues in future work, particularly since some tissues may be more prone to telomere repair than others (Wolf et al., in press). Regardless, our results identify POT1 as an ecologically important gene that may mediate performance in the wild.

Recent work in wild populations shows that telomere elongation can occur amidst a generalized trend of telomere shortening, although it may be limited to specific life-history stages (Fairlie et al., 2016; Hatakeyama et al., 2016; Hoelzl et al., 2016; Spurgin et al., 2018; van Lieshout et al., 2019). Here, we document evidence of short-term telomere elongation in response to acute stress, particularly in chicks of LPS-injected mothers. Our analyses suggest that this elongation is not explained by measurement error (see SI materials, Bateson & Nettle, 2017; Steenstrup et al., 2013). However, alternative mechanisms can manifest as pseudo-lengthening (Epel, 2012), including turnover of existing cells by longer-telomere counterparts or changes in cell composition with age (Beaulieu et al., 2017; Montes et al., 2003; but see Watson et al., 2017). In birds, whole blood is dominated by nucleated red blood cells that turn over every ~9 days (Muriel et al., 2020), and telomere attrition has occurred within a similar timeframe in other species (Nettle et al., 2015; Stier et al., 2016), suggesting that production of longer telomere blood cells is plausible and may co-occur with lower levels of POT1 mRNA. Changes in interstitial telomeres may also appear as elongation, although repeated measures designs like ours minimize these effects (Foote et al., 2013). Moving forward, exploring environmental cues, underlying mechanisms, and long-term consequences of apparent telomere elongation will inform our understanding of telomeres as causal drivers of evolutionarily relevant phenotypes.

Notably, POT1 gene expression varied with our molecular measure of antioxidant capacity. In particular, POT1 and antioxidant gene expression showed a strong positive correlation among chicks, suggesting that individuals may invest in *either* prevention of telomere loss (i.e. high antioxidants but lower telomerase accessibility via high POT1) or recuperation of telomere loss (i.e. higher telomerase accessibility but low antioxidants). Interestingly, male and female chicks may implement different strategies, as 12-day old females exhibited significantly lower POT1 gene expression than males, and some models ≤2 AIC_c_ indicated lower antioxidant gene expression in females as well. POT1 gene expression and change in relative telomere length were the only traits showing significant treatment effects in 12-day old chicks, despite some trends observed for handling-induced CORT and antioxidant gene expression (see **Fig 4**). One interpretation is that these effects are already returning to baseline levels by the time we sampled chicks one week after stress exposure, much like the faded effects of other stressors over time (Deviche et al., 2016; Li et al., 2017). Alternatively, naturalistic stressors like this single day of reduced provisioning may have mild effects on the organism, albeit effects that can culminate in relevant performance consequences (elaborated below). Regardless of these possibilities, our results focus attention on telomere-regulatory mechanisms as potentially vital players in phenotypic responses to early life stress.

### Implications for the evolution of life history strategies

Several components of the chick phenotype also predicted survival to key life history milestones. Telomere dynamics, namely the change in relative telomere length in the week following a stressor and POT1 gene expression, held little importance in predicting immediate survival to fledging; instead, fledging success was higher for heavier chicks and marginally higher for chicks with strong CORT elevation, consistent with past work (McCarty, 2001; Schoenle et al., 2020). On the other hand, change in relative telomere length and POT1 gene expression showed the highest variable importance in predicting recruitment to the breeding population the following year. In particular, chicks with the most positive change in relative telomere length were more likely to return as adults, suggesting that minimized telomere attrition predicts survival (as in Boonekamp et al., 2014; Wood & Young, 2019). This result stands in contrast to most other model comparisons in our study, which showed a higher importance of POT1 gene expression than relative telomere length in predicting fitness-related traits. Notably, though, our sample size of return chicks was quite low, given the intrinsically high juvenile mortality in this system. Considering that the mildest telomere attrition occurred in parallel with downregulation of POT1 gene expression, our results collectively lend strength to the view that underlying telomere regulation may contribute to survival prospects.

Are telomere regulators the driving force connecting telomere dynamics and performance in nature? Telomere regulatory mechanisms do have pleiotropic effects, meaning they may effectively link telomere attrition with downstream performance (Wood & Young, 2019). Indeed, telomerase has dual roles in telomere repair and other vital telomere-independent functions (Ahmed et al., 2008; de Jesus et al., 2011, 2012), yet whether telomerase regulators can equally modulate *both* telomere length and somatic integrity is unclear. As the sole shelterin protein able to bind single-stranded DNA (de Lange, 2018), POT1 is the most direct bridge to telomerase and its downstream effects on telomere length. POT1 also may repress DNA damage (Renfrew et al., 2014; Wu et al., 2020), providing a mechanism for pleiotropic effects that causally link this telomere regulator with more generalized somatic integrity. Under this scenario, telomere length may be a passive scribe accompanying changes in health and longevity (see Bateson & Nettle, 2018), both of which may be driven by ecologically relevant variation in telomere regulators, much like we observed in this study.

### Conclusions

Our results linking POT1 with performance in both adults and chicks suggest that variation in telomere regulators may be visible to natural selection. As molecular biology has infused all areas of ecology, evolution, and behavior, we have repeatedly seen an increasing emphasis on the *regulation* of particular traits, including carotenoids and the maintenance of honesty (Koch et al., 2017; Mundy et al., 2016) or testosterone and the evolution of sexual phenotypes (Fuxjager & Schuppe, 2018; Lipshutz et al., 2019). Our findings advocate for similar changes at the intersection of evolutionary ecology and telomere biology. Clearly, a singular focus on telomere length is incomplete (sensu Casagrande & Hau, 2019; Casagrande et al., 2020; Wood & Young, 2019), and a renewed focus on the mechanisms that shift the balance between attrition and repair is a promising avenue for advances to our understanding of life history.

## ACKNOWLEDGEMENTS

We are grateful to EK Dossey, SD Myers, EM George, DJ Bolinger, BS Duggan, KR Stansberry, and KR Content for support in the field and lab; to the Center for the Integrative Study of Animal Behavior (CISAB) for support; and to our reviewers for feedback. This research was supported by NSF (IOS-1656109; DBI-1460949), the National Institutes of Health (T32HD049336), and the Indiana University Research and Teaching Preserve.

## DATA ACCESIBILITY

The data that support the findings of this study will be openly available on the Dryad Digital Repository.

## AUTHOR CONTRIBUTIONS

SEW and KAR designed the study; SEW and TLS collected samples in the field; SEW coordinated RNA extractions and performed all DNA extractions, qPCR, and statistical analyses; and SEB molecularly sexed all chicks. SEW and KAR drafted the manuscript. All authors read, approved, and contributed to the final manuscript.

## REFERENCES

Ahmed, S., Passos, J. F., Birket, M. J., Beckmann, T., Brings, S., Peters, H.,… Saretzki, G. (2008). Telomerase does not counteract telomere shortening but protects mitochondrial function under oxidative stress. Journal of cell science, 121(7), 1046–1053.

Alatalo, R. V., Lundberg, A., & Glynn, C. (1986). Female pied flycatchers choose territory quality and not male characteristics. Nature, 323(6084), 152.

Allsopp, R. C., Chang, E., Kashefi-Aazam, M., Rogaev, E. I., Piatyszek, M. A., Shay, J. W., & Harley, C. B. (1995). Telomere shortening is associated with cell division in vitro and in vivo. Experimental cell research, 220(1), 194–200.

Angelier, F., Costantini, D., Blevin, P., & Chastel, O. (2018). Do glucocorticoids mediate the link between environmental conditions and telomere dynamics in wild vertebrates? A review. General and comparative endocrinology, 256, 99–111.

Angelier, F., Vleck, C. M., Holberton, R. L., & Marra, P. P. (2015). Bill size correlates with telomere length in male American Redstarts. Journal of Ornithology, 156(2), 525–531.

Angelier, F., Weimerskirch, H., Barbraud, C., & Chastel, O. (2019). Is telomere length a molecular marker of individual quality? Insights from a long◻lived bird. Functional Ecology, 33(6), 1076–1087.

Badas, E. P., Martinez, J., Rivero de Aguilar Cachafeiro, J., Miranda, F., Figuerola, J., & Merino, S. (2015). Ageing and reproduction: antioxidant supplementation alleviates telomere loss in wild birds. J Evol Biol, 28(4), 896–905. doi:10.1111/jeb.12615

Barton, K. (2019). MuMIn: Multi-Model Inference. Retrieved from https://CRAN.R-project.org/package=MuMIn

Bateson, M., & Nettle, D. (2017). The telomere lengthening conundrum–it could be biology. Aging Cell, 16(2), 312–319.

Bauch, C., Becker, P. H., & Verhulst, S. (2013). Telomere length reflects phenotypic quality and costs of reproduction in a long-lived seabird. Proc Biol Sci, 280(1752), 20122540. doi:10.1098/rspb.2012.2540

Bauer, C. M., Graham, J. L., Abolins-Abols, M., Heidinger, B. J., Ketterson, E. D., & Greives, T. J. (2018). Chronological and biological age predict seasonal reproductive timing: an investigation of clutch initiation and telomeres in birds of known age. The American Naturalist, 191(6), 777–782.

Beaulieu, M., Benoit, L., Abaga, S., Kappeler, P. M., & Charpentier, M. J. (2017). Mind the cell: seasonal variation in telomere length mirrors changes in leucocyte profile. Molecular Ecology, 26(20), 5603–5613.

Beaulieu, M., Reichert, S., Le Maho, Y., Ancel, A., & Criscuolo, F. (2011). Oxidative status and telomere length in a long-lived bird facing a costly reproductive event. Functional Ecology, 25(3), 577–585. doi:10.1111/j.1365-2435.2010.01825.x

Beery, A. K., Lin, J., Biddle, J. S., Francis, D. D., Blackburn, E. H., & Epel, E. S. (2012). Chronic stress elevates telomerase activity in rats. Biol Lett, 8(6), 1063–1066. doi:10.1098/rsbl.2012.0747

Benjamini, Y., & Hochberg, Y. (1995). Controlling the false discovery rate: a practical and powerful approach to multiple testing. Journal of the Royal statistical society: series B (Methodological), 57(1), 289–300.

Bentz, A. B., Rusch, D. B., Buechlein, A., & Rosvall, K. A. (2019). The neurogenomic transition from territory establishment to parenting in a territorial female songbird. BMC genomics, 20(1), 1–10.

Blackburn, E. H. (2000). Telomere states and cell fates. Nature, 408(6808), 53–56.

Blackburn, E. H., Greider, C. W., Henderson, E., Lee, M. S., Shampay, J., & Shippen-Lentz, D. (1989). Recognition and elongation of telomeres by telomerase. Genome, 31(2), 553–560.

Bonter, D. N., & Bridge, E. S. (2011). Applications of radio frequency identification (RFID) in ornithological research: a review. Journal of Field Ornithology, 82(1), 1–10.

Boonekamp, J. J., Bauch, C., Mulder, E., & Verhulst, S. (2017). Does oxidative stress shorten telomeres? Biology Letters, 13(5), 20170164.

Boonekamp, J. J., Mulder, G. A., Salomons, H. M., Dijkstra, C., & Verhulst, S. (2014). Nestling telomere shortening, but not telomere length, reflects developmental stress and predicts survival in wild birds. Proceedings of the Royal Society B: Biological Sciences, 281(1785), 20133287. doi:10.1098/rspb.2013.3287

Breuner, C. W., & Berk, S. A. (2019). Using the van Noordwijk and de Jong resource framework to evaluate glucocorticoid-fitness hypotheses. Integrative and Comparative Biology, 59(2), 243–250.

Burnham, K., & Anderson, D. (2002). Model selection and multimodal inference: a practical information-theoretic approach (2nd ed.). New York: Springer.

Burnham, K. P., Anderson, D. R., & Huyvaert, K. P. (2011). AIC model selection and multimodel inference in behavioral ecology: some background, observations, and comparisons. Behavioral Ecology and Sociobiology, 65(1), 23–35.

Çakmak, E., Akın Pekşen, Ç., & Bilgin, C. C. (2017). Comparison of three different primer sets for sexing birds. Journal of Veterinary Diagnostic Investigation, 29(1), 59–63.

Casagrande, S., Stier, A., Monaghan, P., Loveland, J. L., Boner, W., Lupi, S.,… Hau, M. (2020). Increased glucocorticoid concentrations in early life cause mitochondrial inefficiency and short telomeres. Journal of Experimental Biology.

Cawthon, R. M. (2009). Telomere length measurement by a novel monochrome multiplex quantitative PCR method. Nucleic acids research, 37(3), e21–e21.

Chatelain, M., Drobniak, S. M., & Szulkin, M. (2020). The association between stressors and telomeres in non◻human vertebrates: a meta◻analysis. Ecology letters, 23(2), 381–398.

Choi, J., Fauce, S. R., & Effros, R. B. (2008). Reduced telomerase activity in human T lymphocytes exposed to cortisol. Brain, behavior, and immunity, 22(4), 600–605. doi:10.1016/j.bbi.2007.12.004

Costantini, D., Marasco, V., & Moller, A. P. (2011). A meta-analysis of glucocorticoids as modulators of oxidative stress in vertebrates. J Comp Physiol B, 181, 447–456. doi:10.1007/s00360-011-0566-2)

Criscuolo, F., Bize, P., Nasir, L., Metcalfe, N. B., Foote, C. G., Griffiths, K.,… Monaghan, P. (2009). Real-time quantitative PCR assay for measurement of avian telomeres. Journal of Avian Biology, 40(3), 342–347. doi:10.1111/j.1600-048X.2008.04623.x

Criscuolo, F., Smith, S., Zahn, S., Heidinger, B., & Haussmann, M. (2018). Experimental manipulation of telomere length: does it reveal a corner-stone role for telomerase in the natural variability of individual fitness? Phil. Trans. R. Soc. B, 373(1741), 20160440.

Dakin, R., Lendvai, Á. Z., Ouyang, J., Moore, I., & Bonier, F. (2016). Plumage colour is associated with partner parental care in mutually ornamented tree swallows. Animal Behaviour, 111, 111–118.

de Jesus, B. B., Schneeberger, K., Vera, E., Tejera, A., Harley, C. B., & Blasco, M. A. (2011). The telomerase activator TA◻65 elongates short telomeres and increases health span of adult/old mice without increasing cancer incidence. Aging Cell, 10(4), 604–621.

de Jesus, B. B., Vera, E., Schneeberger, K., Tejera, A. M., Ayuso, E., Bosch, F., & Blasco, M. A. (2012). Telomerase gene therapy in adult and old mice delays aging and increases longevity without increasing cancer. EMBO molecular medicine, 4(8), 691–704.

De Lange, T. (2009). How telomeres solve the end-protection problem. Science, 326(5955), 948–952.

de Lange, T. (2018). Shelterin-mediated telomere protection. Annual review of genetics, 52, 223–247.

Deviche, P., Bittner, S., Davies, S., Valle, S., Gao, S., & Carpentier, E. (2016). Endocrine, metabolic, and behavioral effects of and recovery from acute stress in a free-ranging bird. General and comparative endocrinology, 234, 95–102.

Eastwood, J. R., Hall, M. L., Teunissen, N., Kingma, S. A., Hidalgo Aranzamendi, N., Fan, M.,… Peters, A. (2019). Early◻life telomere length predicts lifespan and lifetime reproductive success in a wild bird. Molecular Ecology, 28(5), 1127–1137.

Epel, E. (2012). How “reversible” is telomeric aging? Cancer Prevention Research, 5(10), 1163–1168.

Epel, E. S., Lin, J., Dhabhar, F. S., Wolkowitz, O. M., Puterman, E., Karan, L., & Blackburn, E. H. (2010). Dynamics of telomerase activity in response to acute psychological stress. Brain, behavior, and immunity, 24(4), 531–539.

Fairlie, J., Holland, R., Pilkington, J. G., Pemberton, J. M., Harrington, L., & Nussey, D. H. (2016). Lifelong leukocyte telomere dynamics and survival in a free-living mammal. Aging Cell, 15(1), 140–148. doi:10.1111/acel.12417

Foote, C. G., Vleck, D., & Vleck, C. M. (2013). Extent and variability of interstitial telomeric sequences and their effects on estimates of telomere length. Molecular ecology resources, 13(3), 417–428.

Fox, J., & Weisberg, S. (2011). An {R} Companion to Applied Regression. In: Sage.

Gaunt, A. S., Oring, L. W., Able, K., Anderson, D., Baptista, L., Barlow, J., & Wingfield, J. (1997). Guidelines to the use of wild birds in research. In: Washington, DC: The Ornithological Council.

Geiger, S., Le Vaillant, M., Lebard, T., Reichert, S., Stier, A., Y, L. E. M., & Criscuolo, F. (2012). Catching-up but telomere loss: half-opening the black box of growth and ageing trade-off in wild king penguin chicks. Mol Ecol, 21(6), 1500–1510. doi:10.1111/j.1365-294X.2011.05331.x

George, E. M., & Rosvall, K. A. (2018). Testosterone production and social environment vary with breeding stage in a competitive female songbird. Hormones and behavior, 103, 28–35.

Granger, M. P., Wright, W. E., & Shay, J. W. (2002). Telomerase in cancer and aging. Critical reviews in oncology/hematology, 41(1), 29–40.

Greider, C. W. (1998). Telomerase activity, cell proliferation, and cancer. Proceedings of the National Academy of Sciences, 95(1), 90–92.

Griebel, I. A., Fairhurst, G. D., Marchant, T. A., & Clark, R. G. (2019). Effects of parental and nest-site characteristics on nestling quality in the Tree Swallow (Tachycineta bicolor). Canadian Journal of Zoology, 97(1), 63–71.

Gu, P., Wang, Y., Bisht, K. K., Wu, L., Kukova, L., Smith, E. M.,… Nandakumar, J. (2017). Pot1 OB-fold mutations unleash telomere instability to initiate tumorigenesis. Oncogene, 36(14), 1939–1951.

Hargitai, R., Török, J., Tóth, L., Hegyi, G., Rosivall, B., Szigeti, B., & Szöllősi, E. (2005). Effects of environmental conditions and parental quality on inter-and intraclutch egg-size variation in the Collared Flycatcher (Ficedula albicollis). The Auk, 122(2), 509–522.

Hatakeyama, H., Yamazaki, H., Nakamura, K.-I., Izumiyama-Shimomura, N., Aida, J., Suzuki, H.,… Ishikawa, N. (2016). Telomere attrition and restoration in the normal teleost Oryzias latipes are linked to growth rate and telomerase activity at each life stage. Aging (Albany NY), 8(1), 62.

Haussmann, M. F., Winkler, D. W., Huntington, C. E., Nisbet, I. C., & Vleck, C. M. (2007). Telomerase activity is maintained throughout the lifespan of long-lived birds. Experimental Gerontology, 42(7), 610–618. doi:10.1016/j.exger.2007.03.004

Haywood, S., & Perrins, C. M. (1992). Is clutch size in birds affected by environmental conditions during growth? Proceedings of the Royal Society of London. Series B: Biological Sciences, 249(1325), 195–197.

Hemann, M. T., Strong, M. A., Hao, L.-Y., & Greider, C. W. (2001). The shortest telomere, not average telomere length, is critical for cell viability and chromosome stability. Cell, 107(1), 67–77.

Hoelzl, F., Smith, S., Cornils, J. S., Aydinonat, D., Bieber, C., & Ruf, T. (2016). Telomeres are elongated in older individuals in a hibernating rodent, the edible dormouse (Glis glis). Scientific reports, 6, 36856.

Hussell, D. J. (1983). Age and plumage color in female tree swallows. Journal of Field Ornithology, 312–318.

Hwang, H., Buncher, N., Opresko, P. L., & Myong, S. (2012). POT1-TPP1 regulates telomeric overhang structural dynamics. Structure, 20(11), 1872–1880.

Kawanishi, S., & Oikawa, S. (2004). Mechanism of telomere shortening by oxidative stress. Annals of the New York Academy of Sciences, 1019(1), 278–284.

Kim, S.-Y., & Velando, A. (2015). Antioxidants safeguard telomeres in bold chicks. Biology Letters, 11(5), 20150211.

Koch, R. E., Josefson, C. C., & Hill, G. E. (2017). Mitochondrial function, ornamentation, and immunocompetence. Biological Reviews, 92(3), 1459–1474.

Laprade, H., Querido, E., Smith, M. J., Guérit, D., Crimmins, H., Conomos, D.,… Sfeir, A. (2020). Single-molecule imaging of telomerase RNA reveals a Recruitment-Retention model for telomere elongation. Molecular Cell.

Le Vaillant, M., Viblanc, V. A., Saraux, C., Le Bohec, C., Le Maho, Y., Kato, A.,… Ropert-Coudert, Y. (2015). Telomere length reflects individual quality in free-living adult king penguins. Polar Biology, 38(12), 2059–2067.

Lendvai, A. Z., Akçay, Ç., Weiss, T., Haussmann, M. F., Moore, I. T., & Bonier, F. (2015). Low cost audiovisual playback and recording triggered by radio frequency identification using Raspberry Pi. PeerJ, 3, e877.

Lendvai, A. Z., Ouyang, J. Q., Schoenle, L. A., Fasanello, V., Haussmann, M. F., Bonier, F., & Moore, I. T. (2014). Experimental food restriction reveals individual differences in corticosterone reaction norms with no oxidative costs. PLoS One, 9(11), e110564.

Li, Y., Sun, Y., Krause, J. S., Li, M., Liu, X., Zhu, W.,… Li, D. (2017). Dynamic interactions between corticosterone, corticosteroid binding globulin and testosterone in response to capture stress in male breeding Eurasian tree sparrows. Comparative Biochemistry and Physiology Part A: Molecular & Integrative Physiology, 205, 41–47.

Ludlow, A. T., Gratidao, L., Ludlow, L. W., Spangenburg, E. E., & Roth, S. M. (2017). Acute exercise activates p38 MAPK and increases the expression of telomere◻protective genes in cardiac muscle. Experimental physiology, 102(4), 397–410.

Ludlow, A. T., Lima, L. C., Wang, J., Hanson, E. D., Guth, L. M., Spangenburg, E. E., & Roth, S. M. (2012). Exercise alters mRNA expression of telomere-repeat binding factor 1 in skeletal muscle via p38 MAPK. Journal of Applied Physiology, 113(11), 1737–1746.

Magrath, R. D. (1991). Nestling weight and juvenile survival in the blackbird, Turdus merula. The Journal of Animal Ecology, 335–351.

Martin, T. E., & Geupel, G. R. (1993). Nest-Monitoring Plots: Methods for Locating Nests and Monitoring Success (Métodos para localizar nidos y monitorear el éxito de estos). Journal of Field Ornithology, 507–519.

McCarty, J. P. (2001). Variation in growth of nestling tree swallows across multiple temporal and spatial scales. The Auk, 118(1), 176–190.

McCarty, J. P. (2002). The number of visits to the nest by parents is an accurate measure of food delivered to nestlings in tree swallows. Journal of Field Ornithology, 73(1), 9–14.

Moazzam, M., Yim, T., Kumaresam, V. D., Henderson, D. C., Farrer, L. A., & Zhang, H. (2020). Analysis of telomere length variation and Shelterin complex subunit gene expression changes in ethanol-exposed human embryonic stem cells. Journal of Psychiatric Research.

Monaghan, P., & Ozanne, S. E. (2018). Somatic growth and telomere dynamics in vertebrates: relationships, mechanisms and consequences. Philosophical Transactions of the Royal Society B: Biological Sciences, 373(1741). doi:10.1098/rstb.2016.0446

Montes, I., McLaren, G., Macdonald, D., & Mian, R. (2003). The effects of acute stress on leukocyte activation. J. Physiol. P, 548, 170.

Mundy, N. I., Stapley, J., Bennison, C., Tucker, R., Twyman, H., Kim, K.-W.,… Slate, J. (2016). Red carotenoid coloration in the zebra finch is controlled by a cytochrome P450 gene cluster. Current biology, 26(11), 1435–1440.

Muriel, J., Vida, C., Gil, D., & Pérez-Rodríguez, L. (2020). Ontogeny of leukocyte profiles in a wild altricial passerine. Journal of Comparative Physiology B, 1–12.

Nettle, D., Monaghan, P., Gillespie, R., Brilot, B., Bedford, T., & Bateson, M. (2015). An experimental demonstration that early-life competitive disadvantage accelerates telomere loss. Proc Biol Sci, 282(1798), 20141610. doi:10.1098/rspb.2014.1610

Owen-Ashley, N. T., & Wingfield, J. C. (2006). Seasonal modulation of sickness behavior in free-living northwestern song sparrows (Melospiza melodia morphna). Journal of Experimental Biology, 209(16), 3062–3070.

Palacios, M. G., Winkler, D. W., Klasing, K. C., Hasselquist, D., & Vleck, C. M. (2011). Consequences of immune system aging in nature: a study of immunosenescence costs in free◻living Tree Swallows. Ecology, 92(4), 952–966.

Pineda-Pampliega, J., Herrera-Dueñas, A., Mulder, E., Aguirre, J. I., Höfle, U., & Verhulst, S. (2020). Antioxidant supplementation slows telomere shortening in free-living white stork chicks. Proceedings of the Royal Society B, 287(1918), 20191917.

Pinheiro, J., Bates, D., DebRoy, S., & Sarkar, D. (2019). nlme: Linear and Nonlinear Mixed Effects Models. Retrieved from https://CRAN.R-project.org/package=nlme

Pisoschi, A. M., & Pop, A. (2015). The role of antioxidants in the chemistry of oxidative stress: A review. European journal of medicinal chemistry, 97, 55–74.

Reichert, S., & Stier, A. (2017). Does oxidative stress shorten telomeres in vivo? A review. Biology Letters, 13(12), 20170463.

Renfrew, K. B., Song, X., Lee, J. R., Arora, A., & Shippen, D. E. (2014). POT1a and components of CST engage telomerase and regulate its activity in Arabidopsis. PLoS Genet, 10(10), e1004738.

Rosvall, K. A., Reichard, D. G., Ferguson, S. M., Whittaker, D. J., & Ketterson, E. D. (2012). Robust behavioral effects of song playback in the absence of testosterone or corticosterone release. Hormones and behavior, 62(4), 418–425.

Schoenle, L. A., Zimmer, C., Miller, E. T., & Vitousek, M. N. (2020). Does variation in glucocorticoid concentrations predict fitness? A phylogenetic meta-analysis. General and comparative endocrinology, 113611.

Schwagmeyer, P., & Mock, D. W. (2008). Parental provisioning and offspring fitness: size matters. Animal Behaviour, 75(1), 291–298.

Spurgin, L. G., Bebbington, K., Fairfield, E. A., Hammers, M., Komdeur, J., Burke, T.,… Richardson, D. S. (2018). Spatio◻temporal variation in lifelong telomere dynamics in a long◻term ecological study. Journal of Animal Ecology, 87(1), 187–198.

Sridhar, M., Thammiah, V., & Suganthi, R. (2014). Detection of changes in genes related to antioxidant and biotransformation function in broiler birds fed aflatoxin by real time PCR. The Indian Journal of Animal Sciences, 84(2).

Steenstrup, T., Hjelmborg, J. v. B., Kark, J. D., Christensen, K., & Aviv, A. (2013). The telomere lengthening conundrum—artifact or biology? Nucleic acids research, 41(13), e131–e131.

Stier, A., Delestrade, A., Bize, P., Zahn, S., Criscuolo, F., & Massemin, S. (2016). Investigating how telomere dynamics, growth and life history covary along an elevation gradient in two passerine species. Journal of Avian Biology, 47(1), 134–140.

Stutchbury, B. J., & Robertson, R. J. (1986). A simple trap for catching birds in nest boxes. Journal of Field Ornithology, 57(1), 64–65.

Stutchbury, B. J., & Robertson, R. J. (1987). Two Methods of Sexing Adult Tree Swallows before They Begin Breeding (Dos Metodos para Determinar el Sexo en Golondrinas (Tachycineta bicolor)). Journal of Field Ornithology, 236–242.

Sudyka, J. (2019). Does Reproduction Shorten Telomeres? Towards Integrating Individual Quality with Life◻History Strategies in Telomere Biology. Bioessays.

Tinbergen, J., & Boerlijst, M. (1990). Nestling weight and survival in individual great tits (Parus major). The Journal of Animal Ecology, 1113–1127.

Vagasi, C. I., Pătra◻, L., Pap, P. L., Vincze, O., Mure◻an, C., Nemeth, J., & Lendvai, A. Z. (2018). Experimental increase in baseline corticosterone level reduces oxidative damage and enhances innate immune response. PLoS One, 13(2), e0192701.

van Lieshout, S. H., Bretman, A., Newman, C., Buesching, C. D., Macdonald, D. W., & Dugdale, H. L. (2019). Individual variation in early◻life telomere length and survival in a wild mammal. Molecular Ecology, 28(18), 4152–4165.

Verhulst, S., Aviv, A., Benetos, A., Berenson, G. S., & Kark, J. D. (2013). Do leukocyte telomere length dynamics depend on baseline telomere length? An analysis that corrects for ‘regression to the mean’. European journal of epidemiology, 28(11), 859–866.

Virgin, E. E., & Rosvall, K. A. (2018). Endocrine-immune signaling as a predictor of survival: A prospective study in developing songbird chicks. General and comparative endocrinology, 267, 193–201.

Vitousek, M. N., Taff, C. C., Ardia, D. R., Stedman, J. M., Zimmer, C., Salzman, T. C., & Winkler, D. W. (2018). The lingering impact of stress: brief acute glucocorticoid exposure has sustained, dose-dependent effects on reproduction. Proceedings of the Royal Society B: Biological Sciences, 285(1882), 20180722.

von Zglinicki, T. (2002). Oxidative stress shortens telomeres. Trends in Biochemical Sciences, 27(7), 339–344.

Wada, H., Hahn, T. P., & Breuner, C. W. (2007). Development of stress reactivity in white-crowned sparrow nestlings: total corticosterone response increases with age, while free corticosterone response remains low. General and comparative endocrinology, 150(3), 405–413.

Wang, F., Podell, E. R., Zaug, A. J., Yang, Y., Baciu, P., Cech, T. R., & Lei, M. (2007). The POT1–TPP1 telomere complex is a telomerase processivity factor. Nature, 445(7127), 506.

Watson, R. L., Bird, E. J., Underwood, S., Wilbourn, R. V., Fairlie, J., Watt, K.,… McNeilly, T. N. (2017). Sex differences in leucocyte telomere length in a free◻living mammal. Molecular Ecology, 26(12), 3230–3240.

Wei, T., Simko, V., Levy, M., Xie, Y., Jin, Y., & Zemla, J. (2017). Package ‘corrplot’. Statistician, 56, 316–324.

Wilbourn, R. V., Moatt, J. P., Froy, H., Walling, C. A., Nussey, D. H., & Boonekamp, J. J. (2018). The relationship between telomere length and mortality risk in non-model vertebrate systems: a meta-analysis. Philosophical Transactions of the Royal Society B: Biological Sciences, 373(1741), 20160447.

Winkler, D. W., Hallinger, K. K., Pegan, T. M., Taff, C. C., Verhoeven, M. A., Chang Van Oordt, D.,… Andersen, M. J. (2020). Full lifetime perspectives on the costs and benefits of lay date variation in tree swallows. Ecology.

Wood, E. M., & Young, A. J. (2019). Telomere attrition predicts reduced survival in a wild social bird, but short telomeres do not. Molecular Ecology.

Wu, Y., Poulos, R. C., & Reddel, R. R. (2020). Role of POT1 in Human Cancer. Cancers, 12(10), 2739.

Yarru, L. P., Settivari, R. S., Gowda, N. K. S., Antoniou, E., Ledoux, D. R., & Rottinghaus, G. E. (2009). Effects of turmeric (Curcuma longa) on the expression of hepatic genes associated with biotransformation, antioxidant, and immune systems in broiler chicks fed aflatoxin. Poultry science, 88(12), 2620–2627. doi:10.3382/ps.2009-00204

Zakian, V. A. (2012). Telomeres: the beginnings and ends of eukaryotic chromosomes. Experimental cell research, 318(12), 1456–1460.

Zietzer, A., Buschmann, E., Janke, D., Li, L., Brix, M., Meyborg, H.,… Hillmeister, P. (2017). Acute physical exercise and long◻term individual shear rate therapy increase telomerase activity in human peripheral blood mononuclear cells. Acta Physiologica, 220(2), 251–262.

Zinzow-Kramer, W. M., Horton, B. M., & Maney, D. L. (2014). Evaluation of reference genes for quantitative real-time PCR in the brain, pituitary, and gonads of songbirds. Horm Behav, 66(2), 267–275. doi:10.1016/j.yhbeh.2014.04.011

