## Supplementary material for "The telomere regulatory gene POT1 responds to stress and predicts performance in nature: implications for telomeres and life history evolution": SI materials

**Method S1. RFID Methodology**

Nest boxes were equipped with radio-frequency identification (RFID) readers. At each box, we installed a circular coil antenna (28G copper magnet wire coated in truck bed liner or electrical tape, 7.5cm outer/6.5cm inner diameter, inductance ~125mH) around the entrance of the nest box. Antennae were connected to a “Generation 2” reader (Bonter & Bridge, 2011), which was powered by a 12V battery and housed inside a waterproof junction box below the nest box (described in Lendvai, Akçay, Weiss, et al., 2015). Birds were outfitted with passive integrated transponder (PIT) tags (2.3 mm; EM4102, IB Technology, Leicester, UK), and reader detection of PIT tags cycled every 400ms, with 300ms signal detection and 100ms pause, continuously from 06:00 to 21:00 EDT, to save battery life. RFID readers recorded a unique PIT tag ID and time stamp every time a PIT tag passed through the antennae, and therefore, the entrance to the box.

On days 4, 5, and 6 post-hatching, readers recorded all PIT tag IDs detected at the box entrance, including any banded parents. We classified the two highest-read PIT tags at each nest box as the male and female parents. All highest-read female PIT tags correctly matched the female injected during the manipulation, and banded males caught during the experiment were also correctly matched. All but 2 females were equipped with PIT tags prior to injection. In 8 nests, RFID readers yielded 0 reads for at least one day of the experiment, which indicates that either the RFID reader, antennae, or PIT tags were not working properly, and were therefore excluded from the analysis.

We used RFID boards during the chick period to measure visitation rates. We determined the number of nest box visits by first filtering out continuous readings occurring within 3 seconds of another read of the same individual, often generated when a bird is perched at the nest entrance. To account for entrances and exits, we then estimated the number of visits by an individual as half the number of remaining reads. Visitation rate could overestimate the actual number of feeding visits (e.g. approaching antennae but not entering or exiting); however, previous studies in tree swallows show that feeding occurs during the majority (95-96%) of visits to the nest (McCarty, 2002; Whittingham et al., 2003), suggesting that overall, RFID estimates of feeding are accurate.

This method has previously been validated in tree swallows (Lendvai et al., 2015) and other bird species using similar calculations (Farine et al., 2012; García-Navas et al., 2009; Nomano et al., 2014), where visitation rates estimated from RFID and personal observations were highly correlated (r=0.69-0.99). We also validated the accuracy of RFID logs in this study, where we observed 30-min baseline (n=16) and post-injection (n=35) visitation rates during the afternoon prior to injection or 3-6h after injection. Post-injection observers were blind to treatment. During observations, we recorded the number of female/male entrances and exits, times perched at the hole, and the total number of visits to the box, as sex could not always be distinguished. Later, we used RFID logs to calculate visitation at the same date and time as personal observations, in order to ask whether RFID estimates are indicative of visitation rates. We found a significant positive correlation between observed and RFID estimated visitation rates (F_1,53_=6.14, p=0.0168, n=55, r=0.903).

Females and males share in provisioning, and so we opportunistically evaluated any indirect effects of female treatment on male partners. Because males are harder to capture early in the breeding season, only 18/43 males in this study had PIT tags. Nevertheless, we analyzed male visitation (n=15 males with usable data) and found no difference in visitation rate between the male partners of saline and LPS-injected females during the presumed “peak” of LPS-induced sickness (i.e. 3-6 hours post-injection, F_1,13_=0.09, p=0.91). This result matches a previous study by Palacios et al. (2011), showing that male tree swallows did not compensate for LPS-induced decreases in female visitation rates. In fact, other work indicates that males might mirror female decreases in parental care, e.g. when a female partner is handicapped via wing clipping, as in Winkler and Allen (1995).

**Method S2. Maternal Injections**

Our LPS injection protocol followed Palacios et al. (2011), using a subcutaneous injection of either saline or lipopolysaccharide (LPS) saline-oil emulsion in the right dorsal apterium (n=22 saline, 21 LPS). We injected each bird with 50µL of 0.2mg/mL LPS solution (0.5mg/kg body weight, based on an average female mass of 20.05g). The LPS saline-oil emulsion consisted of LPS from *Escherichia coli* (serotype 055:B5, lot #086M4146V; Sigma-Aldrich, St. Louis, Missouri, USA) dissolved in 0.9% sterile saline. We then emulsified the solution at a 1:1 ratio with Freund’s incomplete adjuvant (Sigma-Aldrich), which prolongs the expression of sickness symptoms up to 48h (Owen-Ashley & Wingfield, 2006). A previous study using this method in breeding female tree swallows reported body mass reduction, lethargy, and decreased parental behaviors in LPS-treated birds (Palacios et al., 2011). We injected all control birds with a saline-oil emulsion.

**Method S3. Chick Sexing Protocol**

We used the automated Maxwell® RSC Instrument (Promega, Madison, WI) and Whole Blood DNA Kit (#AS1520) to extract DNA from ≤25µL red blood cells, which was eluted into 60-70µL buffer. We determined the sex of each chick following established methods (Çakmak et al., 2017). Each reaction contained 5µL PerfeCTA SYBR Green SuperMix No Rox (Quanta Biosciences), 0.2µL 200nM P2 primer, 0.2µL 200nM P8 primer (see Table S1), 3.6µL dH_2_O, and 1µL extracted DNA (range: 100-300 ng/µL). We ran PCR under the following conditions: (95°/2m) + 40x (95°/30s, 51°/45s, 72°/45s) + 72°/5m + 4° hold. 8µL of each PCR product was mixed with 1.5µL loading dye and run on a 3% EtBr agarose gel (1.5h at 90V) alongside a known adult male and female sample. Males exhibited a single band at ~250bp and females exhibited a double band at ~250 and 275bp. All chicks were successfully sexed. As validation, we also ran DNA from breeding adults (n=32 per sex), which exhibited a brood patch/cloacal protuberance or were raising chicks with another known-sex individual, and gel results were accurate with a 0.05% error rate.

**Method S4. Validation of Telomere Lengthening**

When examining the distribution of telomere changes from pre-treatment to 12-days old, we observed some increases in telomere length over time within individuals. We conducted a sensitivity analysis to compare the scope of this change to the scope of technical variation in qPCR. We estimated differences in telomere lengths among technical replicates, i.e., triplicates next to each other within a plate, and we compared these technical replicates to variation in biological replicates, i.e., difference in RTL from pre-treatment to 12-days old within an individual chick. We used MCMglmm (Hadfield, 2010) with an inverse Wishart prior (*v* = 1, nu = 0.002), 600,000 iterations, a thinning of 300 and burn-in period of 15,000 iterations, to test whether within-individual changes in RTL were greater than measurement error. We used two samples per individual and built a model with RTL as the response variable and individual ID and plate ID as random effects (*n* = 294 samples, 147 individuals). We then randomly selected one set of triplicates per individual and constructed a model with RTL for each of the technical replicates as the response variable and individual ID as a random effect (*n* = 147 individuals). We compared the variance explained by the random effect for individual ID between these two models and whether the 95% confidence intervals overlapped. Additionally, we separated the data set into groups with positive changes in relative telomere length (n = 72 individuals) or negative changes in relative telomere length (n = 75 individuals), and we ran these models again for these groups separately, as in van Lieshout et al. (2019).

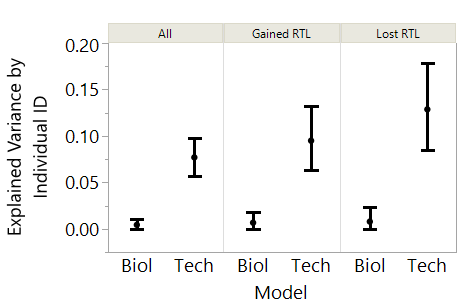
Increases in RTL were identified in 48.9% of within-individual changes (n = 72). When accounting for plate effects using MCMCglmm, the random effect estimate for individual ID with technical replicates was 0.077 (95% CI = 0.057, 0.098), whereas for within-individual samples the random effect estimate was 0.0044 (95% CI = 0.0001, 0.01). For the group that exhibited increases in RTL the random effect estimate for individual ID with technical replicates was 0.095 (95% CI = 0.063, 0.13), whereas for within-individual samples this estimate was 0.0067 (95% CI = 0.0002, 0.018). The random effect estimate for technical replicates in the group that exhibited decreases in RTL was 0.13 (95% CI = 0.085, 0.18) and for within-individual samples this estimate was 0.0079 (0.0002, 0.023). None of the 95% CI’s from the technical replicates and within-individual samples overlapped (**Fig S1**).

**Fig S1.** A sensitivity analysis showing that the random effect estimate of individual ID explained more variance in relative telomere length among technical replicates (Tech) than the change in biological replicates (Biol) from pre-treatment to 12-days old, indicating that increases in relative telomere length with age were not due to measurement error alone.

| **Table S1.** Primer sequences for gene expression analyses | |
| --- | --- |
| **Gene Name** | **Primer Sequence (5’ to 3’)** |
| Peptidylprolyl isomerase A (PPIA) forward | TCCCGAAGACAGCAGAAAACT |
| Peptidylprolyl isomerase A (PPIA) reverse | CCATTGTGGCGTGTGAAGTC |
| Superoxide dismutase (SOD) forward | CAGGATTCTGTCATTTCCCTTTC |
| Superoxide dismutase (SOD) reverse | CGTTTCCAGTTAACTTGCTCTC |
| Peroxiredoxin-1 (PRDX-1) forward | GCTTCTGTTGACTCTCACTTCT |
| Peroxiredoxin-1 (PRDX-1) reverse | CCTTCAGCACTCCATACTCTTT |
| Glutathione Peroxidase (GPX) forward | CGCAGTACATCATCTGGTCTC |
| Glutathione Peroxidase (GPX) reverse | GTCCTGGATCTTGATGGTTTCA |
| Protection of telomeres 1 (POT1) forward | CCTCGACAGTATCGCATTAGAG |
| Protection of telomeres 1 (POT1) reverse | GCAGAGCCACGTAGAATGAA |
| Glyceraldehyde 3-phosphate dehydrogenase (GAPDH) forward | ACCAGCCAAGTACGATGACAT |
| Glyceraldehyde 3-phosphate dehydrogenase (GAPDH) reverse | CCATCAGCAGCAGCCTTCA |
| Telomere (Telc) forward | ACACTAAGGTTTGGGTTTGGGTTTGGGTTTGGGTTAGTGT |
| Telomere (Telg) reverse | GTTAGGTATCCCTATCCCTATCCCTATCCCTATCCCTAACA |
| P2 (sexing primer) | TCTGCATCGCTAAATCCTTT |
| P8 (sexing primer) | CTCCCAAGGATGAGRAAYTG |

| **Table S2.** The top models (∆AIC_c_ ≤ 2) predicting treatment effects on chick physiology. The global models included treatment, sex, treatment x sex interaction, brood size, and hatch date, and random effect of nest. qPCR plate ID was included for change in relative telomere length (∆RTL), time of day included for CORT measurements. K = # of parameters (including intercept), w_i_ = model weight, R^2^_m_ = variance explained by fixed effect, R^2^_c_ = variance explained by fixed and random effects. g.e. = gene expression. | | | | | | | |
| --- | --- | --- | --- | --- | --- | --- | --- |
| **Variable** | **Model** | **K** | **AICc** | **∆AIC_c_** | **w_i_** | **R^2^_m_** | **R^2^_c_** |
| Growth (n=159) | Treatment*Sex + Brood Size | 5 | 1158.66 | 0 | 0.296 | 0.198 | 0.599 |
|  | Treatment*Sex + Brood + Hatch Date | 6 | 1159.4 | 0.74 | 0.205 | 0.232 | 0.603 |
|  | Treatment + Sex + Brood Size | 4 | 1160.52 | 1.86 | 0.117 | 0.199 | 0.601 |
| ∆ RTL (n=147) | Treatment + qPCR Plate | 3 | 41.80 | 0 | 0.792 | 0.042 | 0.167 |
| POT1 g.e. (n=77) | Treatment + Sex | 3 | 207.79 | 0 | 0.508 | 0.161 | 0.339 |
|  | Treatment*Sex | 4 | 209.67 | 1.88 | 0.198 | 0.163 | 0.348 |
| PC1 antioxidant g.e. (n=77) | Treatment*Sex | 4 | 291.44 | 0 | 0.309 | 0.1 | 0.221 |
|  | Treatment + Sex | 3 | 291.46 | 0.02 | 0.306 | 0.09 | 0.185 |
|  | Treatment | 2 | 293.09 | 1.65 | 0.135 | 0.036 | 0.19 |
| Baseline CORT (n=107) | Treatment + Time of Day | 3 | 259.95 | 0 | 0.362 | 0.011 | 0.074 |
|  | Treatment | 2 | 261.1 | 1.15 | 0.204 | 0.004 | 0.069 |
| Handling- CORT (n=107) | Treatment + Time of Day | 3 | 244.56 | 0 | 0.476 | 0.06 | 0.278 |

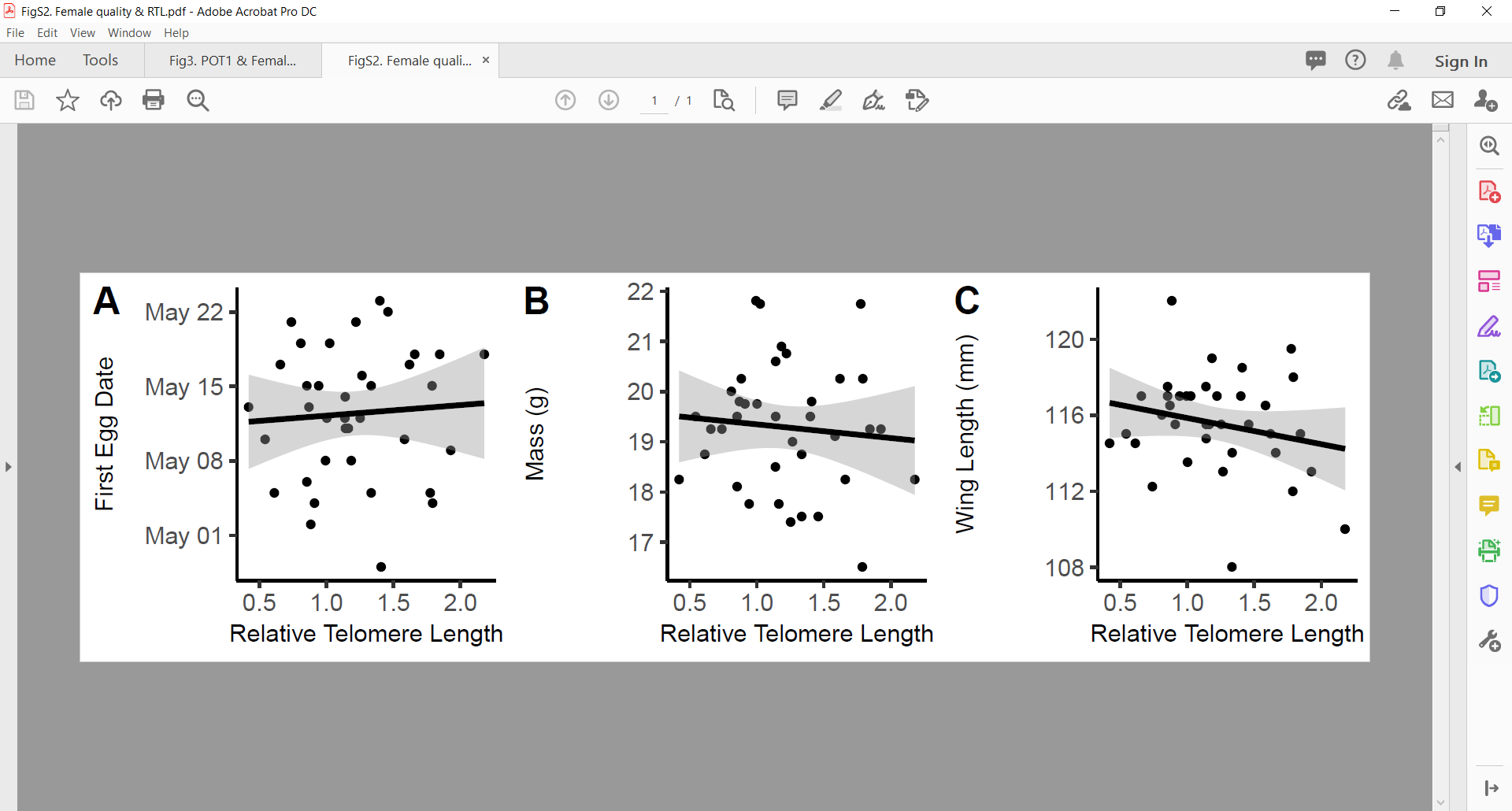
**Fig S2.** Relative telomere length does not predict female (A) first egg date, (B) body mass, or (C) wing length. Shading indicates 95% confidence intervals from model output.

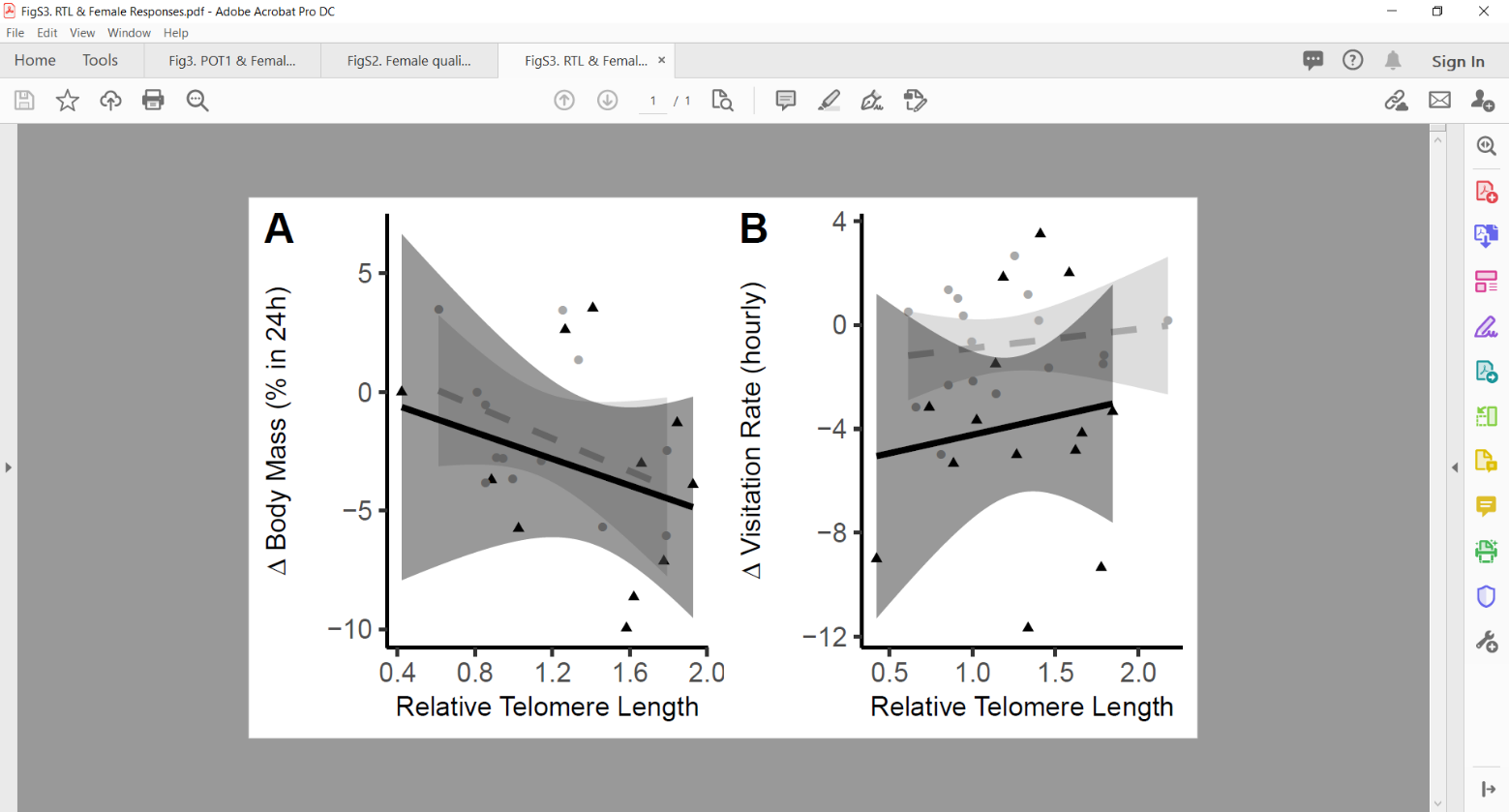
**Fig S3**. Female relative telomere length and responses to stress following injections of breeding mothers with saline (gray, circles, dashed line) or LPS (black, triangles, solid line): A) female ∆ body mass within 24h of injection or B) ∆ visitation rate during the peak of sickness.

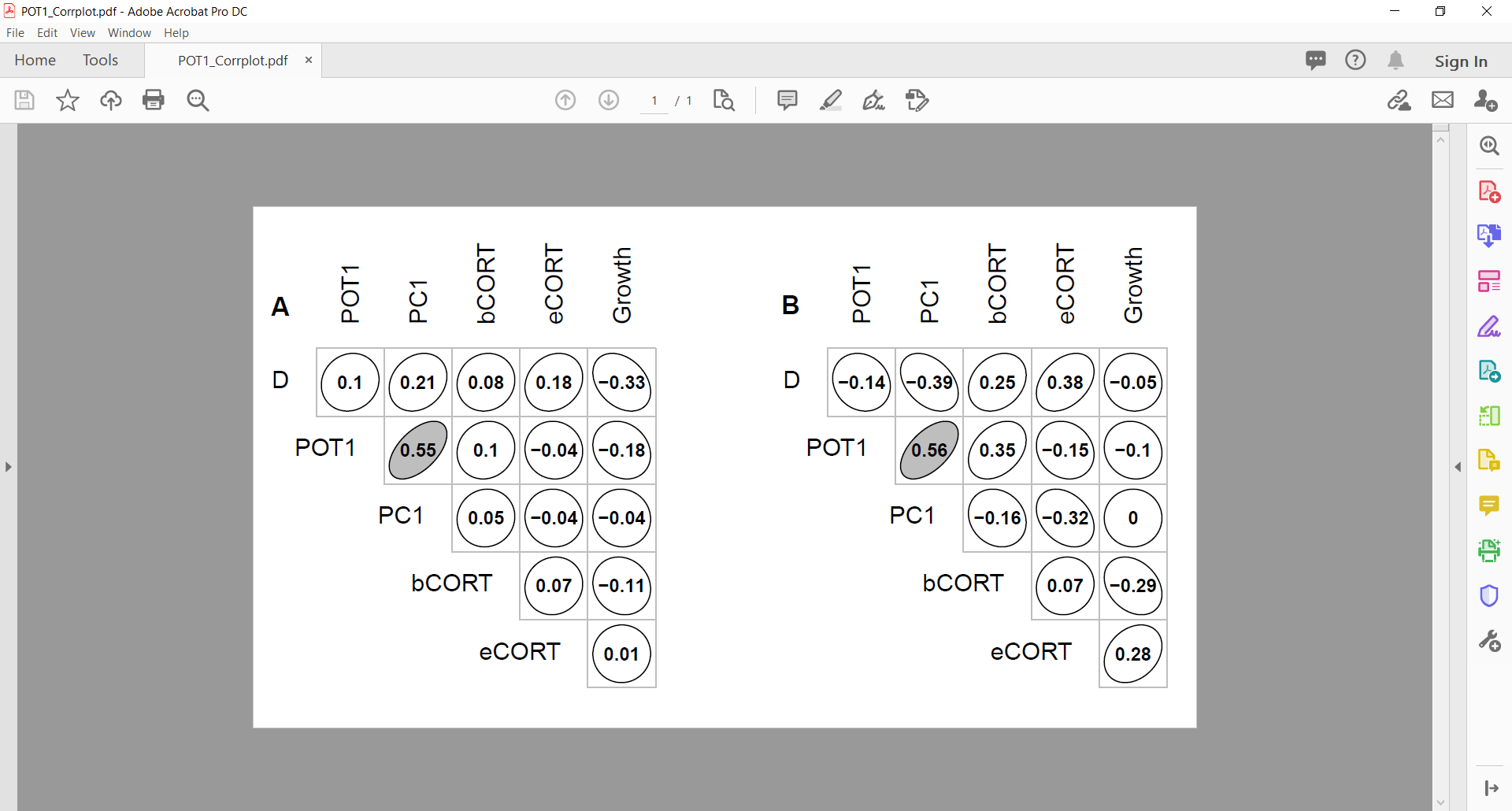

**Fig S4.** Spearman’s correlation values between different chick phenotypes. Ellipse shape denotes the strength and direction of correlations in chicks of saline (A) and LPS-injected (B) mothers. Significant correlations are shaded in gray.

**References**

Bonter, D. N., & Bridge, E. S. (2011). Applications of radio frequency identification (RFID) in ornithological research: a review. *Journal of Field Ornithology, 82*(1), 1-10.

Çakmak, E., Akın Pekşen, Ç., & Bilgin, C. C. (2017). Comparison of three different primer sets for sexing birds. *Journal of Veterinary Diagnostic Investigation, 29*(1), 59-63.

Farine, D. R., Garroway, C. J., & Sheldon, B. C. (2012). Social network analysis of mixed-species flocks: exploring the structure and evolution of interspecific social behaviour. *Animal Behaviour, 84*(5), 1271-1277.

García-Navas, V., Ortego, J., & Sanz, J. J. (2009). Heterozygosity-based assortative mating in blue tits (Cyanistes caeruleus): implications for the evolution of mate choice. *Proceedings of the Royal Society of London B: Biological Sciences, 276*(1669), 2931-2940.

Lendvai, A. Z., Akçay, Ç., Ouyang, J. Q., Dakin, R., Domalik, A. D., St John, P. S., . . . Bonier, F. (2015). Analysis of the optimal duration of behavioral observations based on an automated continuous monitoring system in tree swallows (Tachycineta bicolor): is one hour good enough? *PLoS One, 10*(11), e0141194.

Lendvai, A. Z., Akçay, Ç., Weiss, T., Haussmann, M. F., Moore, I. T., & Bonier, F. (2015). Low cost audiovisual playback and recording triggered by radio frequency identification using Raspberry Pi. *PeerJ, 3*, e877.

McCarty, J. P. (2002). The number of visits to the nest by parents is an accurate measure of food delivered to nestlings in tree swallows. *Journal of Field Ornithology, 73*(1), 9-14.

Nomano, F. Y., Browning, L. E., Nakagawa, S., Griffith, S. C., & Russell, A. F. (2014). Validation of an automated data collection method for quantifying social networks in collective behaviours. *Behavioral Ecology and Sociobiology, 68*(8), 1379-1391.

Owen-Ashley, N. T., & Wingfield, J. C. (2006). Seasonal modulation of sickness behavior in free-living northwestern song sparrows (Melospiza melodia morphna). *Journal of Experimental Biology, 209*(16), 3062-3070.

Palacios, M. G., Winkler, D. W., Klasing, K. C., Hasselquist, D., & Vleck, C. M. (2011). Consequences of immune system aging in nature: a study of immunosenescence costs in free‐living Tree Swallows. *Ecology, 92*(4), 952-966.

van Lieshout, S. H., Bretman, A., Newman, C., Buesching, C. D., Macdonald, D. W., & Dugdale, H. L. (2019). Individual variation in early‐life telomere length and survival in a wild mammal. *Molecular Ecology, 28*(18), 4152-4165.

Whittingham, L. A., Dunn, P. O., & Clotfelter, E. D. (2003). Parental allocation of food to nestling tree swallows: the influence of nestling behaviour, sex and paternity. *Animal Behaviour, 65*(6), 1203-1210.

Winkler, D. W., & Allen, P. E. (1995). Effects of handicapping on female condition and reproduction in tree swallows (Tachycineta bicolor). *The Auk, 112*(3), 737-747.
